# A new tractable method for generating Human Alveolar Macrophage Like cells in vitro to study lung inflammatory processes and diseases

**DOI:** 10.1101/2023.04.05.535806

**Authors:** Susanta Pahari, Eusondia Arnett, Jan Simper, Abul Azad, Israel Guerrero-Arguero, Chengjin Ye, Hao Zhang, Hong Cai, Yufeng Wang, Zhao Lai, Natalie Jarvis, Miranda Lumbreras, Diego Jose Maselli-Caceres, Jay Peters, Jordi B Torrelles, Luis Martinez-Sobrido, Larry S Schlesinger

## Abstract

Alveolar macrophages (AMs) are unique lung resident cells that contact airborne pathogens and environmental particulates. The contribution of human AMs (HAM) to pulmonary diseases remains poorly understood due to difficulty in accessing them from human donors and their rapid phenotypic change during *in vitro* culture. Thus, there remains an unmet need for cost-effective methods for generating and/or differentiating primary cells into a HAM phenotype, particularly important for translational and clinical studies. We developed cell culture conditions that mimic the lung alveolar environment in humans using lung lipids, *i.e.*, Infasurf (calfactant, natural bovine surfactant) and lung-associated cytokines (GM-CSF, TGF-β, and IL-10) that facilitate the conversion of blood-obtained monocytes to an AM-Like (AML) phenotype and function in tissue culture. Similar to HAM, AML cells are particularly susceptible to both *Mycobacterium tuberculosis* and severe acute respiratory syndrome coronavirus 2 (SARS-CoV-2) infections. This study reveals the importance of alveolar space components in the development and maintenance of HAM phenotype and function, and provides a readily accessible model to study HAM in infectious and inflammatory disease processes, as well as therapies and vaccines.

**IMPORTANCE:** Millions die annually from respiratory disorders. Lower respiratory track gas-exchanging alveoli maintain a precarious balance between fighting invaders and minimizing tissue damage. Key players herein are resident AMs. However, there are no easily accessible *in vitro* models of HAMs, presenting a huge scientific challenge. Here we present a novel model for generating AML cells based on differentiating blood monocytes in a defined lung component cocktail. This model is non-invasive, significantly less costly than performing a bronchoalveolar lavage, yields more AML cells than HAMs per donor and retains their phenotype in culture. We have applied this model to early studies of *M. tuberculosis* and SARS-CoV-2. This model will significantly advance respiratory biology research.

## INTRODUCTION

Alveolar macrophages (AMs) live in a unique tissue environment and must maintain lung homeostasis through recycling of alveolar lining fluid and surfactant lipids, as well as clearance of inhaled debris and microbes without damaging the alveoli and impairing gas exchange (1). AMs’ importance in maintaining lung homeostasis is evident in individuals with pulmonary alveolar proteinosis (PAP) where AM development and function are impaired, resulting in the accumulation of pulmonary surfactant that obstructs the airways (2). AMs can self-maintain in the steady state (3) and also originate from peripheral blood monocytes and fetal monocytes (4, 5). AM functions are regulated by alveolar type-II epithelial cells through their interactions with CD200 and transforming growth factor-β (TGF-β) leading to IL-10 secretion, which is important for cell homeostasis (1). TGF-β itself is important for AM development (6). Granulocyte macrophage colony-stimulating factor (GM-CSF), secreted by resident macrophages and lung epithelial cells, is also essential for AM development (7, 8). Generation of a non-transformed, GM-CSF– dependent murine macrophage line shows some similarity with mouse AMs (9). GM-CSF induces the transcription factor peroxisome proliferator-activated receptor gamma (PPAR-γ), which is highly expressed by AMs and critical for AM development (7).

It is increasingly appreciated that tissue environments greatly influence macrophage phenotype and function (10, 11) and that AMs are distinct from other macrophages including lung interstitial macrophages (IMs). For example, AMs are less glycolytic than IMs and highly express genes involved in oxidative phosphorylation (OxPhos) and fatty acid metabolism (12). AMs also respond to stimuli differently than other tissue macrophages. The lung surfactant protein (SP)-A specifically increases mannose receptor (MR/CD206; MRC1, a signature of AMs) expression in AMs, but not in peritoneal macrophages (13) and also drives IL-4-mediated AM proliferation and activation in the lung, but not in the peritoneal cavity (14). AMs are also more susceptible to infection by the intracellular pathogen *Mycobacterium tuberculosis* (*M.tb*) than other tissue macrophages, including IMs in the lung (12). PPAR-γ enhances *M.tb* growth specifically in lung macrophages, but not in bone marrow-derived or peritoneal macrophages (15). In addition, *M.tb* infection of AMs *in vivo* is distinct from infection of AMs that have been out of the lung for 18 h (16), likely because the transcriptome of AMs rapidly changes after removal from the lung (16, 17). The unique nature of AMs and their loss of phenotype after removal from the lung make the study of AM biology and the impact of AMs on infectious and non-infectious diseases challenging.

There are no current tractable and easily accessible *in vitro* models of human AMs (HAM). One method of acquiring HAM is by bronchoalveolar lavage (BAL), which is expensive, invasive, labor intensive (18) and only recovers ∼2-4 x 10^6^ HAM per person. This is particularly problematic during the Coronavirus Disease 2019 (COVID-19) pandemic, which has limited many research procedures, including performing human BALs, thus making it even harder to study HAM biology. Another method is to obtain HAM from cadaveric lung tissue of recently deceased individuals, which is accessible to only a few labs. Murine AMs are relatively more easily obtainable, but BAL results in only ∼3-5x 10^5^ AMs per mouse (19) and cellular pathways of interest may deviate from those found in HAM.

Since transplanting peritoneal macrophages into the lung results in loss of peritoneal markers and gain of PPAR-γ and other AM markers (10), we hypothesized that culturing human monocytes in lung components would drive them to an AM-Like (AML) phenotype, thus providing a more readily available model to study HAM. AMs constantly ingest and catabolize surfactant lipids that line the alveoli and are exposed to locally produced cytokines such as GM-CSF, TGF-β and IL-10. Thus, we developed an AML cell model by culturing readily available human blood-derived monocytes [in peripheral blood mononuclear cells (PBMCs) or purified] with an optimized lung component cocktail composed of GM-CSF, TGF-β, IL-10 and Infasurf, a natural bovine-derived surfactant replacement therapeutic that contains phospholipids [26 mg phosphatidylcholine (PC) with 16 mg as desaturated PC], neutral lipids like cholesterol, and 0.7 mg hydrophobic SP-B and SP-C. Infasurf does not contain SP-A and SP-D. Our initial optimization study demonstrated that both SP-A and SP-D are not important for AM differentiation and development. Indeed, we focused on core elements that are more constant for human cell AM development among donors than the other components of alveolar lining fluid.

Infasurf, GM-CSF, TGF-β and IL-10 signaling resulted in upregulation of PPAR-γ, a signature transcription factor essential for AM development. Human AML cells exhibited light and electron microscopy morphology resembling HAM, including the appearance of lipid body inclusions, some appearing as lamellar bodies. Expression of a gene set unique to HAM as well as global transcriptomic analysis by RNA-seq revealed expression profiles of AML cells related to freshly obtained HAM, including increased expression of key AM transcription factors and PPAR-γ, TGF-β and GM-CSF signaling pathways. In addition, AML cells showed increased OxPhos and mitochondrial respiration and reduced glycolysis, similar to what is reported for AMs (12). AML cells had increased expression of CD206, macrophage receptor with collagenous structure (MARCO) and CD11c and reduced CD36 expression. Culturing AML macrophages in the lung component cocktail after macrophage adherence maintained the AML phenotype over time in culture. Importantly, similar to HAM, AML cells were particularly susceptible to the airborne pathogens, *Mycobacterium tuberculosis* (*M.tb)* and SARS-CoV-2. Thus, we present a novel model for generating AML cells which is minimally-invasive, significantly less costly, results in more AML cells relative to HAM recovered from one person and can be maintained in culture. Individual components of the cocktail alone cannot generate AML cells. We present a promising model to study HAM in a variety of lung inflammation contexts.

## RESULTS

### *In vitro* development and differentiation of human AML cells

We established a method for providing exogenous surfactant components (Infasurf) and specific lung-associated cytokines (GM-CSF, TGF-β, IL-10), critical for AM differentiation, to cultured monocytes in peripheral blood mononuclear cells (PBMCs) to determine whether the lung-associated components would drive monocyte differentiation into macrophages resembling a HAM phenotype (20). Importantly, we analyzed freshly obtained HAM within 6h of acquisition which best enables retention of the *in vivo* phenotype (17).

We isolated PBMCs from healthy adult human donors and first cultured them in increasing concentrations of GM-CSF, TGF-β and IL-10 without Infasurf for 6 days, during which time monocytes differentiated into macrophages. We identified the optimal concentration of these cytokines to induce expression of PPARG and MRC1, two well-established AM markers **(Fig. S1A-B)**. Next, to understand the role of individual lung-associated cytokines and surfactant in generating AML cells, we treated PBMCs with GM-CSF, TGF-β, IL-10 and Infasurf individually or in combination and assessed expression of a subset of genes that are differentially expressed in HAM relative to MDM. Treatment with all four components (termed “ALL cocktail”) drove more robust gene expression changes than individual cytokines or Infasurf treatment alone **(Fig. S1C-K).** ALL cocktail treatment did not affect the viability of AML cells (**Fig. S1L**). Next, we cultured PBMCs in the presence of ALL cocktail at the optimal concentration for 6 days (ALL cocktail) **(Fig. 1A).** To assess the cultured macrophages further, we identified a set of 30 genes that are differentially expressed in fresh HAM compared with blood-based monocyte-derived macrophages (MDM) **(Table 1)**. These genes were chosen carefully based on the literature and a previously generated AmpliSeq database from our lab comparing MDM and HAM transcriptomes (17). We assessed the gene expression pattern in the cultured macrophages from a randomly selected subset of these genes from **Table 1** that resemble the HAM phenotype. Monocytes cultured in ALL cocktail developed into macrophages that exhibited expression patterns similar to HAM with significant increases in expression of PPARG, MRC1, MARCO, CES1, MCEMP1, MCL1, DUSP1, CXCL3, PU.1, CXCL5, CD170 and CCL18 and significant decreases in expression of MMP7, MMP9, CD36, CCL22 and CD84 when compared to monocytes that were cultured without lung components and thus differentiated into MDM **(Fig. 1B-Q)**. We named the cells cultured in ALL cocktail AM-Like (AML) cells. Increases in PPARG transcript in AML cells *vs.* MDM corresponded with an increase in PPARG protein levels **(Fig 1R)**. The transcription factor PU.1 (SPI1) is induced by GM-CSF and is important for AM function (8). Like HAM, both AML and MDM expressed PU.1, although increased in AML cells **(Fig 1J, R)**. Thus, the established culture conditions drive both PPARG and PU.1 expression, critical transcriptional determinants of AML development.

**Fig 1.**
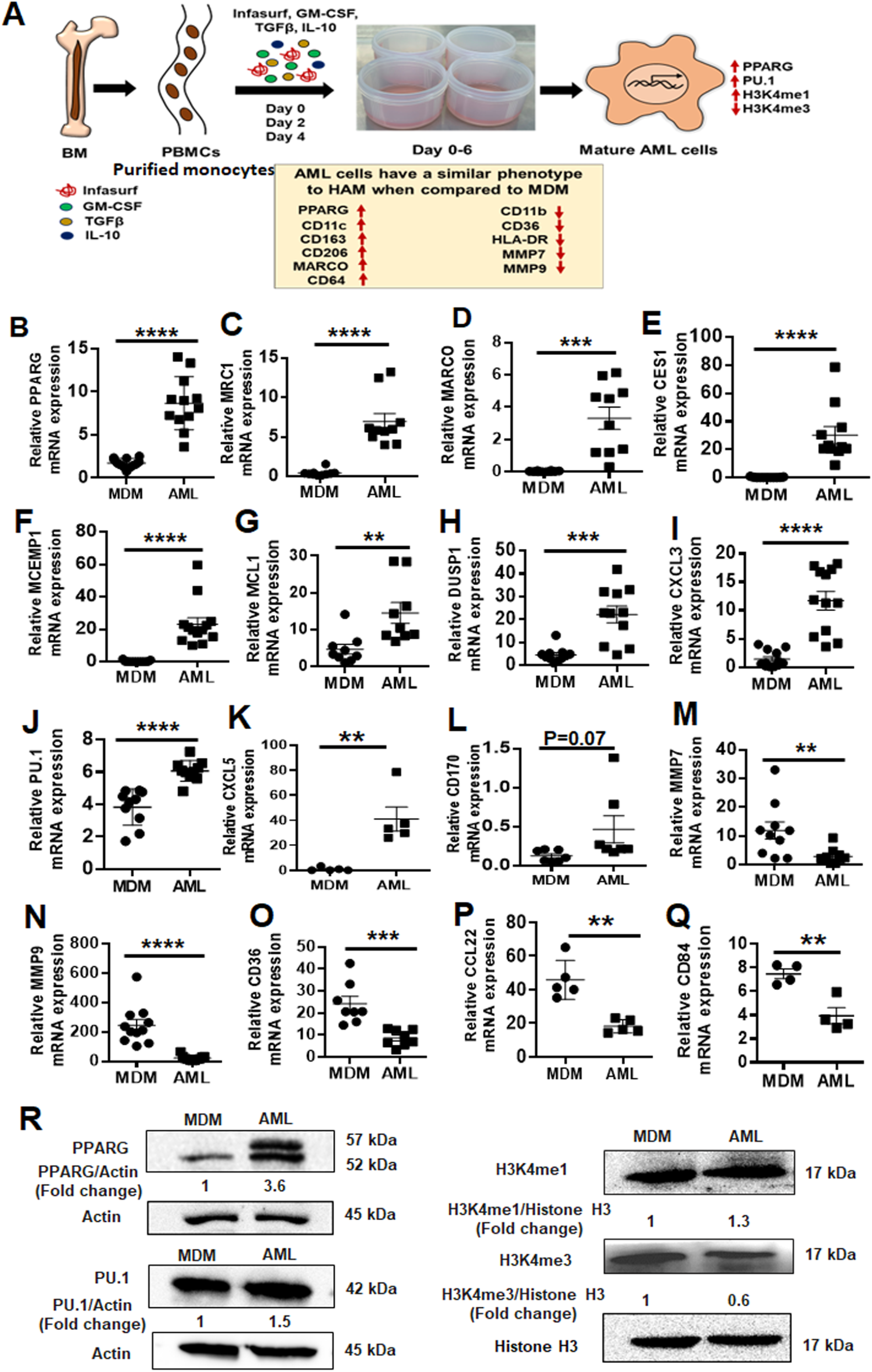
AML cells exhibit a similar phenotype to HAM when compared to MDM. (A) Model of *in vitro* generation of human Alveolar Macrophage-Like (AML) cells from human PBMCs. Healthy human PBMCs were exposed (Day 0, 2, 4) to lung-associated components [surfactant (Infasurf) and cytokines (GM-CSF, TGF-β, IL-10)] (“ALL cocktail”) for 6 days or left untreated (MDM). AML cells demonstrated a similar phenotype to HAM (17, 21) compared to MDM with indicated higher (red upside arrow) and lower (red downside arrow) cell surface expression. AML cells and HAM have similar transcriptional profiles with increased expression of PPARG and PU.1 (SPI1). Like HAM (17, 21), AML cells express specific histone modifications and methylation with high H3K4me1 and low H3K4me3. (B-Q) PBMCs were exposed to ALL cocktail for 6 days on alternative days (Day 0, 2, 4) or left untreated (MDM). qRT-PCR data demonstrate significant increases in (B) PPARG, (C) MRC1, (D) MARCO, (E) CES1, (F) MCEMP1, (G) MCL1, (H) DUSP1, (I) CXCL3, (J) PU.1 (SPI1), (K) CXCL5 and (L) CD170 and decreases in (M) MMP7, (N) MMP9, (O) CD36, (P) CCL22 and (Q) CD84 expression in AML cells compared to untreated MDM. Gene expression was normalized to Actin. Representative dot plots showing relative mRNA expression of the indicated genes from 12-15 human donors. Each dot indicates individual donors. Data are expressed as mean ± SEM and analyzed by Unpaired Student’s ‘t’ test ** p≤0.01, ***p≤0.001, ****p≤0.0001. (R) AML cells demonstrate a HAM-like phenotype, with increased expression of PPARG, PU.1, H3K4me1 and decreased expression of H3K4me3. Nuclear extracts were collected and Western blot performed to assess expression of PPARG, PU.1, H3K4me1 and H3K4me3. Actin and Histone H3 were used as loading controls. Representative blots from n=4, numbers below each blot indicate mean fold change relative to MDM.

**Table 1.**
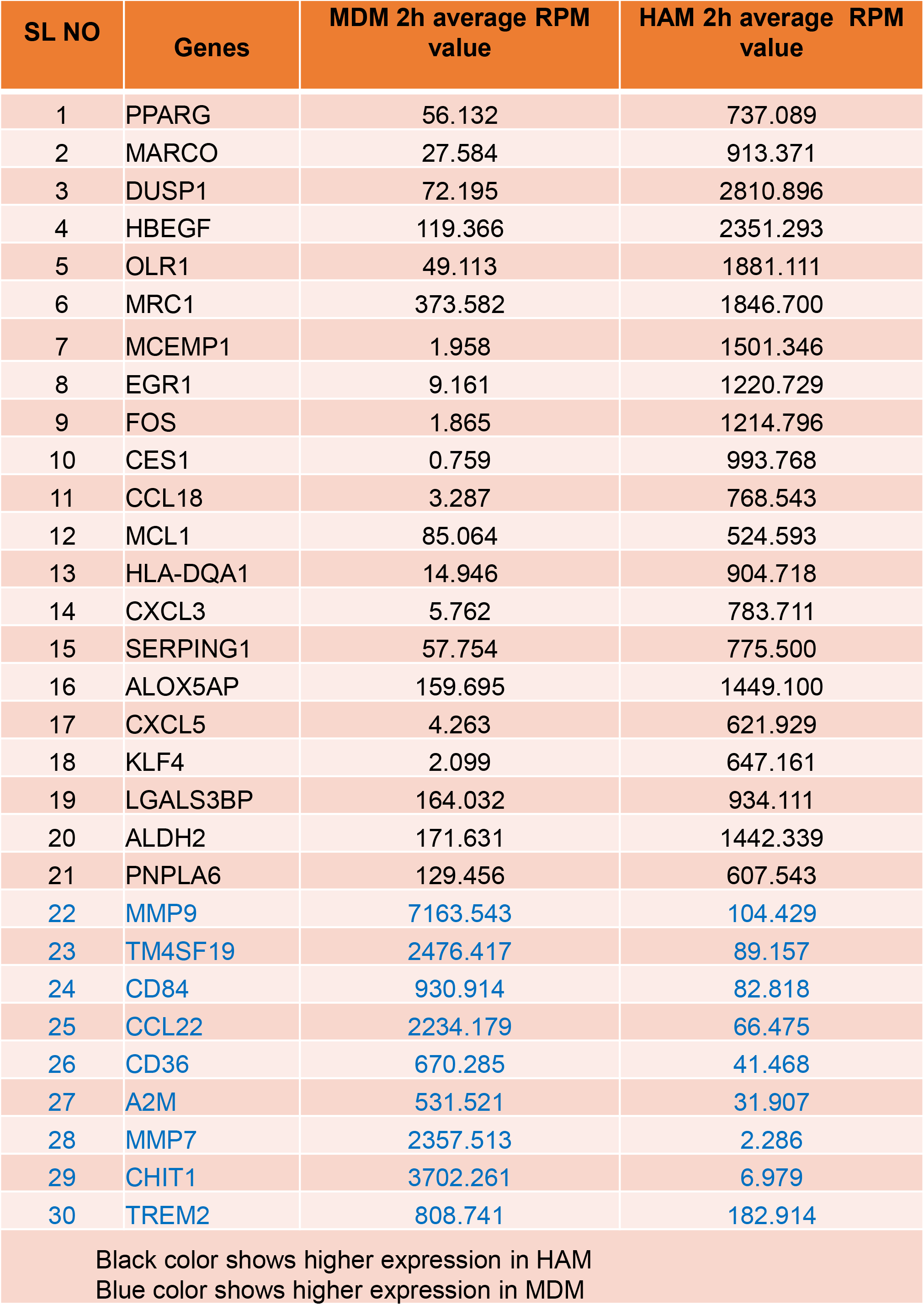
Thirty gene signature to differentiate MDM and HAM. **Gene list to differentiate MDMs and HAMs** These genes were carefully chosen based on their abundancy in HAM relative to MDM, from the literature and a previously generated AmpliSeq database from our laboratory (17, 20). Transcriptomic analysis was assessed from PBMC-derived differentiated MDM and fresh HAM after 2h adherence in culture. ‘RPM’ means ‘reads per million’, i.e., the count of how many reads map to the gene, divided by the total number of aligned reads and multiplied by one million.

### AML cells undergo similar epigenetic changes as reported for HAM

During development, AMs undergo specific histone modifications, with higher levels of histone H3 lysine 4 mono-methylation (H3K4me1) and lower levels of H3K4me3 (21). These epigenetic changes result in the recruitment of PU.1, which is essential for maintenance of high H3K4me1 at macrophage-specific enhancers (22). As observed for HAM (21), AML cells also showed higher expression of H3K4me1 and lower expression of H3K4me3 when compared to MDM **(Fig. 1R)** (20). These data indicate that culturing human monocytes in lung components during differentiation drives them to an AML phenotype with characteristics similar to HAMs.

### Continuous supplementation of the lung cocktail retains the AM phenotype after differentiation of AML cells

To determine whether continuous supplementation of the ALL cocktail during monocyte differentiation is necessary to drive monocytes to AML cells, we treated PBMCs with one dose of ALL cocktail (Day 0) *vs.* multiple doses (Days 0, 2, 4). Monocytes treated with ALL cocktail on alternative days showed changes in gene expression more akin to HAM when compared to one dose only (Day 0). We observed a stronger increase in PPARG, MRC1, MARCO, CES1, PU.1 and MCEMP1 gene and protein expression when cells were treated with multiple cocktail doses **(Fig. 2A-G)**. Together, the results indicate that monocytes must be continuously supplemented with ALL cocktail during differentiation to drive the monocytes to AML cells.

**Fig 2.**
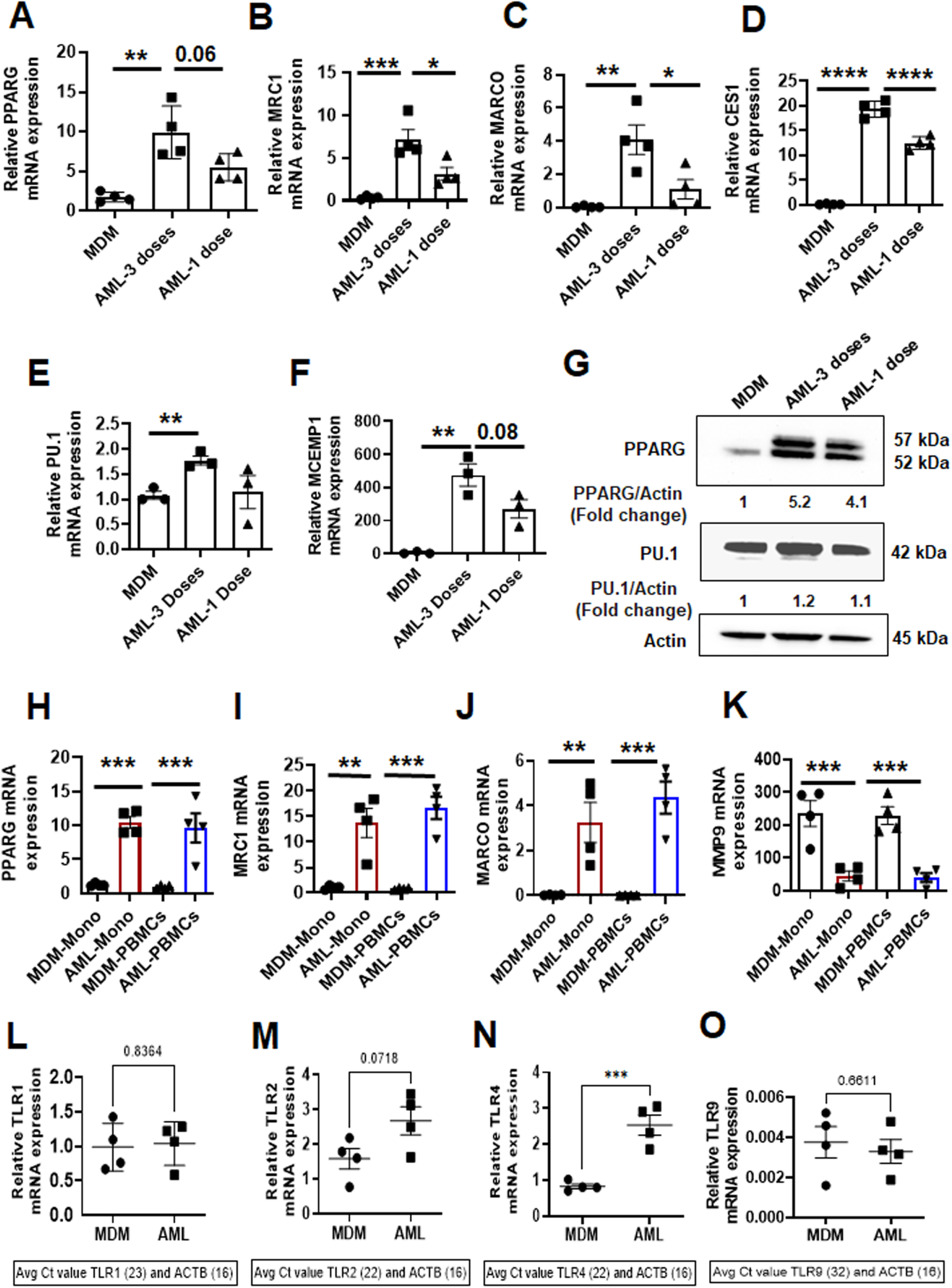
Continuous supplementation of the lung component cocktail during differentiation is necessary to drive monocytes to AML cells. (A-F) PBMCs from healthy human donors were exposed to ALL cocktail [surfactant (Infasurf: 100 µg/mL) and cytokines (GM-CSF: 10 ng/mL, TGF-β: 5 ng/mL, IL-10: 5 ng/mL)] for 6 days after only one administration on Day 0 (1 dose), on alternative days (3 doses), or left untreated (MDM). Gene expression of (A) PPARG, (B) MRC1, (C) MARCO, (D) CES1, (E) PU.1 and (F) MCEMP1 was significantly higher in AML cells that received 3 doses of treatment than 1 or 0 dose. Each dot indicates an individual donor, n=4. (G) PPARG and PU.1 protein levels were also higher in AML cells stimulated with all 3 doses. Actin was used as loading control. Representative blots from 2 human donors, the numbers below each blot indicate mean fold change relative to MDM. (H-K) Monocytes were purified by EasySep™ human monocyte isolation kit from healthy human PBMCs on Day 0 (Mono) and exposed to ALL cocktail [AML-Mono: surfactant (Infasurf: 100 µg/mL) and cytokines (GM-CSF: 10 ng/mL, TGF-β: 5 ng/mL, IL-10: 5 ng/mL)] for 6 days on alternative days or left untreated (MDM-Mono). In addition, PBMCs were exposed to ALL cocktail for 6 days on alternative days (AML-PBMCs) or left untreated (MDM-PBMCs), then macrophages were purified by adherence. The cells were collected for qRT-PCR analysis of selected HAM signature genes (17). Gene expression was compared within the groups: MDM and AML cells that were matured from purified monocytes (MDM-Mono and AML-Mono) and MDM and AML cells that were matured in the PBMCs (MDM-PBMCs and AML-PBMCs), then macrophages were isolated. The qRT-PCR data show gene expression of (H) PPARG, (I) MRC1, (J) MARCO and (K) MMP9 expressed as relative mRNA expression normalized to Beta actin control. Each dot indicates an individual donor. Data are expressed as mean ± SEM (n=4) and analysed by Ordinary one-way ANOVA with Sidak’s multiple comparisons test. *p≤0.05, ** p≤0.01, ***p≤0.001, ****p≤0.0001. Differential expression of relevant TLR genes in AML cells and MDM are shown in (L) TLR1, (M) TLR2, (N) TLR4 and (O) TLR9. n=4. Each dot indicates an individual donor. Data are expressed as mean ± SEM and analyzed by Unpaired Student’s ‘t’ test *** p≤0.001.

To determine if the lymphocytes present in PBMCs aid in differentiation of monocytes into AML cells, we assessed expression of select genes for AML cells generated from PBMCs *vs.* purified monocytes during cultivation. We found that lymphocytes are not required for AML cell development. AML cells generated from PBMCs and isolated monocytes showed similar increases in expression of PPARG, MRC1 and MARCO, and reduced expression of MMP9 **(Fig. 2H-K)**. We also observed significant differential expression of TLR genes in AML cells vs. MDM **(Fig. 2L-O)**. These data indicate that AML cells can be developed from monocytes in the absence or presence of other cell types present in PBMCs. For ease, unless indicated otherwise we cultured PBMCs with 3 doses of ALL cocktail to generate the AML cells described below.

We have previously determined (AC Papp, 2018. S1 Table) that AMs rapidly lose their phenotype upon isolation from the lung and time in culture (17). Our data demonstrate that treatment with multiple doses of ALL cocktail (Days 0, 2, 4) is optimal for AML development **(Fig. 2A-G).** To investigate if continuous supplementation of the cocktail is required to retain the AM phenotype after differentiation of AML cells, we generated AML cells, plated them and subsequently incubated them with or without ALL cocktail for 24h, 48h and 72h. We observed that additional supplementation of cocktail after adherence enables maintenance of the AML phenotype with higher expression of PPARG, MRC1 and MARCO compared to cells that are not treated after adherence **(Fig. S2A-C)**. This phenotype can be maintained for a longer period of time with ALL cocktail added **(Fig. S3A-C)**, which is beneficial for longer-term studies.

### AML cells have similar morphological features and limited self-proliferation capacity to those reported for HAM

HAM have a unique morphology (23). We assessed the morphology of MDM and AML cells by light microscopy and transmission electron microscopy (TEM). By light microscopy AML cells were more rounded and had long pseudopodia closely resembling HAMs, as opposed to MDMs which are flatter, more irregularly elongated cells **(Fig 3A-B)**. Similarly, by TEM, AML cells appeared rounded and had a similar morphology to what is reported for HAM (23). Like HAM, the cytoplasm of AML cells contained various structures which vary in appearance and number **(Fig 3C)**. AML cells contained prominent onion shaped phagolysosomes with phospholipid rich surfactant stored in lipid inclusion bodies, resembling lamellar bodies (LB), composite bodies (CB), and large and small floccular or reticular inclusions, some showing fusion (23). We also observed coated vesicles, some large heterophagic vacuoles (HV), Palade granules (PG), and very dense granules interpreted as ferritin (F) (23) **(Fig 3C)**. Double membrane autophagosome (DMA) structures were also visible as were several round/irregular or elongated mitochondria (M) and various elements of the endoplasmic reticulum (ER). In contrast to AML cells or HAM, MDM were flat, large, and irregularly shaped with an eccentrically placed nucleus, numerous vesicles and vacuoles, a ruffled surface, and contained free or membrane-bound lysosomal inclusions in the vacuole. Round or ovoid electron-dense bodies (EDB) were more prevalent in MDM than AML cells. Round or elongated mitochondrial structures and ER were also visible in MDM **(Fig 3D)**. Overall, the micrographs showed that AML cells have morphology similar to that described for HAM, which represent morphologically distinct macrophages (23).

**Fig 3.**
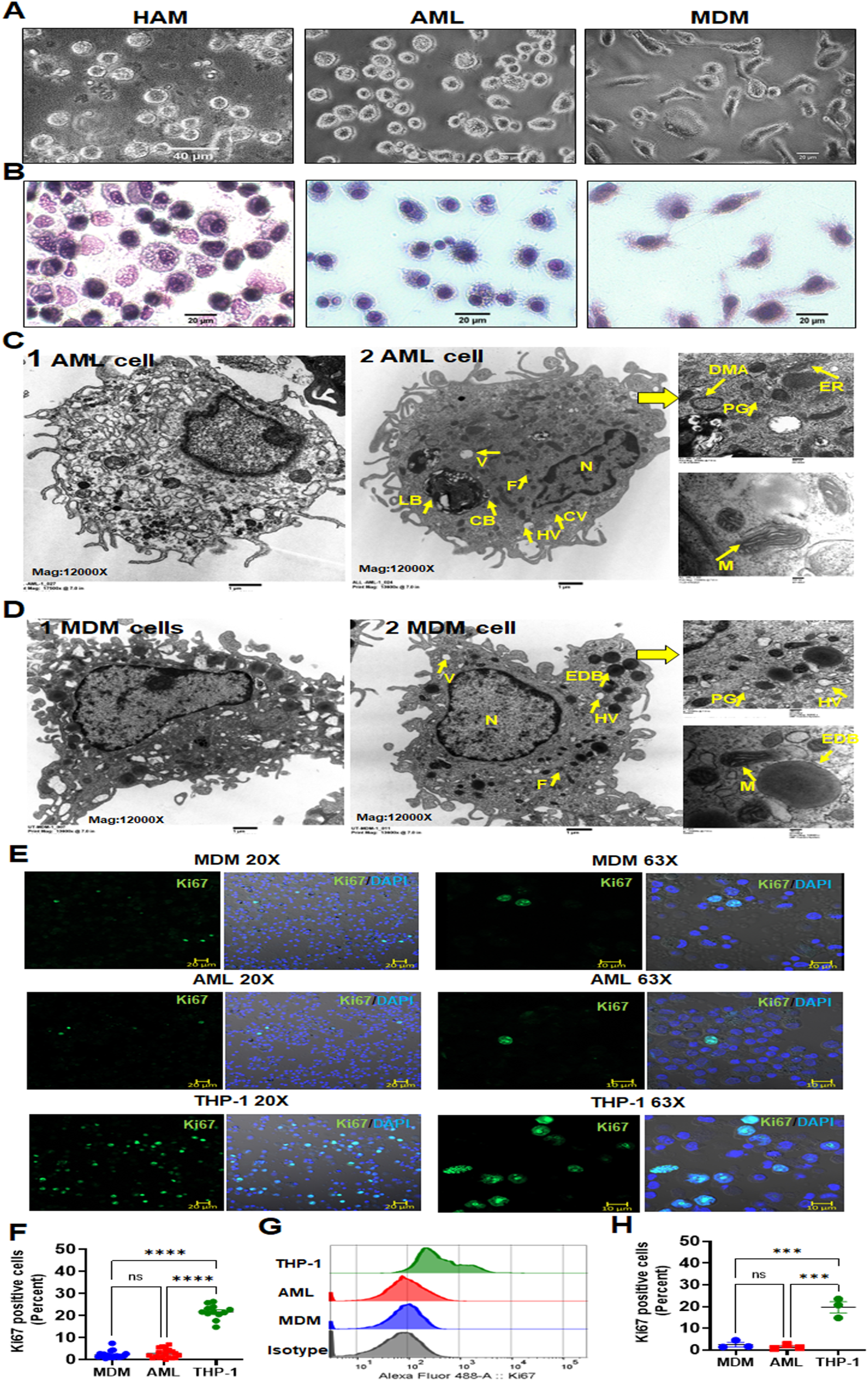
AML cells are morphologically similar to HAM when compared to MDM. (A) Light microscopy images of HAM, AML and MDM cells indicating that AML cells have a more rounded appearance resembling HAM. (B) Morphology of AML cells was compared with HAM and MDM after cytospin and staining with HEMA 3 by light microscopy. (C, D) Representative TEM images (1 & 2) of AML cells and MDM, scale bar: 1 µm. AML cells are rounded with long pseudopodia similar to what has been reported using TEM on HAM from healthy adult human donors (23). (C) AML cells contain onion shaped phago(lyso)somes with phospho lipid-rich surfactant material stored in lipid inclusion bodies, named as lamellar bodies [LB], composite bodies [CB], coated vesicles [CV], heterophagic vacuoles [HV], double membrane autophagosomes [DMA], round/irregular or elongated mitochondria [M], palade granules [PG], ferritin [F], endoplasmic reticulum [ER], nucleus [N]. (D) MDM are irregularly shaped with an eccentrically placed nucleus [N], ER, numerous vesicles [CV] and vacuoles [V], and ruffled surface, free or membrane-bound lysosomal inclusions in the vacuole. Round or ovoid electron-dense bodies [EDB], Palade granules [PG], Ferritin (F), round or elongated mitochondria [M] are more abundant in MDM. (C-D) Magnification: 12,000x, Higher magnification insets on the right: 50,000x, Scale bar: 200 nm. (E, F) MDM, AML and THP-1 monocytic cells were immunostained with Ki67 antibody (green) and DAPI for nucleus (blue), then imaged with confocal microscopy. Scale bar: 10µm, 20µm and 20x, 63x magnification. (F) Confocal data of Ki67 positive cells (percent) were quantified from >200 macrophages (DAPI positive cells) per microscopic field. Each dot indicates a separate field. Cumulative data from 3 donors, mean ± SEM and analyzed with one-way ANOVA. ****p≤0.0001. (G, H) Flow cytometry histogram data (G) show representative Ki67 MFI and (H) each dot indicates percent positive cells, n=3 donors. Data are expressed as mean ± SEM with one-way ANOVA. ***p = <0.001, ns=non-significant.

Adult AMs have been described as long-lived terminally differentiated lung resident cells (24). In the murine model, AMs originate from either peripheral blood monocytes and/or fetal monocytes, and undergo cell proliferation for self-renewal and maintenance in the steady state (4, 5, 25, 26). In contrast, much less is known about the ontogeny and cell proliferation of healthy adult human AMs in the normal steady state condition. Some evidence suggests a low-grade proliferation capacity of adult human AMs in disease states such as respiratory infection or inflammatory/autoimmune diseases but not in healthy humans (27, 28). To ascertain the proliferative capacity of AML cells and MDM, cells were stained using Ki67 and analyzed by confocal microscopy and flow cytometry **(Fig. 3E-H)**. AML cells and MDM demonstrated very limited proliferation capacity. The THP-1 monocytic cell line was used as a positive control.

### HAM and AML cells share similar transcriptome profiles

To further compare AML cells to HAM and MDM, we performed transcriptomic analysis following RNA-seq of freshly collected HAM, AML cells and MDM. Principal component analysis (PCA) showed a high degree of similarity between biological replicates within each group **(Fig. 4A)**. Volcano plots depicting the false discovery rate (FDR) relative to the magnitude of change in gene expression highlighted that the majority of the AML cell transcriptomes resemble HAM, with 899 genes (of 14,097 expressed genes; 6.4%) significantly upregulated at least 2-fold, and 102 genes (0.7%) significantly down regulated at least 2-fold in AML cells relative to HAM **(Fig. 4B, D).** In contrast, when comparing MDM and freshly isolated HAM, we found a significant difference with 1,516 up-regulated, and 1,319 down-regulated genes in MDM *vs.* HAM **(Fig. 4C,** 20.1% genes were differentially expressed). We also found differential gene expression with 744 up-regulated and 438 down-regulated genes in AML cells when compared to the MDM transcriptome **(Fig. S4A)**. Results from the RNA-seq data validated the 30 gene signature comparing fresh HAM to MDM **(Table 1; Fig. 1; Fig. 4E, F)**.

**Fig 4.**
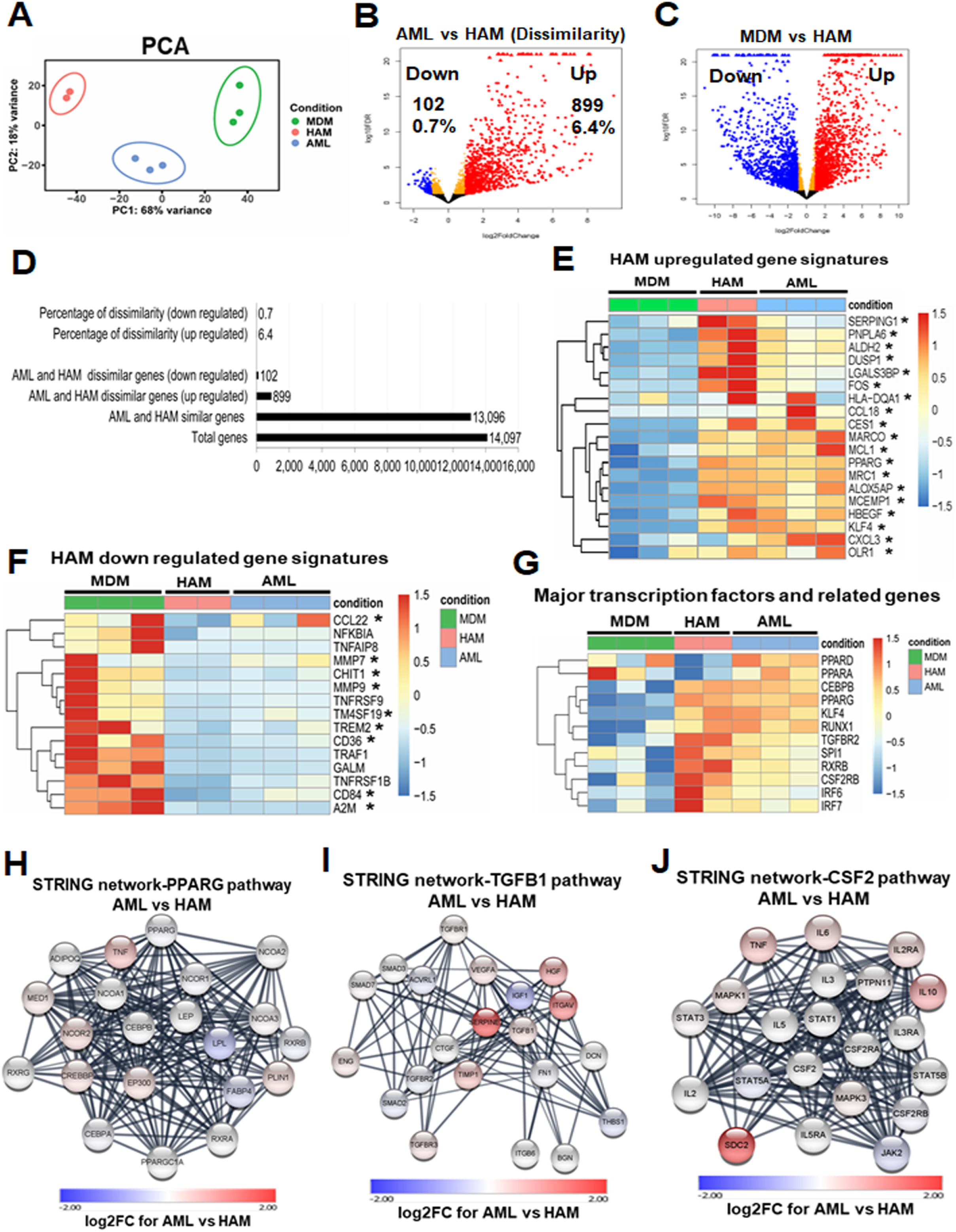
HAM and AML cells share similar transcriptional profiles and related pathways. (A) Principal-component analysis (PCA) demonstrates minimal variation within the biological replicates (HAM: n=2 donors; AML: n=3 donors; MDM: n=3 donors). (B) Volcano plot demonstrates the comparison between the AML and HAM transcriptome. AML and HAM are similar: out of 14,097 expressed genes, only 899 genes are up-regulated ≥ 2-fold with FDR adjusted p-value < 0.05 (red), and 102 genes are down-regulated (blue) in AML cells. (C) Volcano plot demonstrates the comparison between MDM and HAM transcriptome. MDM and HAM are more dissimilar: out of 14,097 expressed genes, 1,516 are up-regulated (red) and 1,319 are down-regulated (blue) in MDM. (D) Bar graph represents the comparison between the AML and HAM transcriptome. (E, F) Heatmaps showing major up- and down-regulated genes in MDM, HAM and AML cells. The asterisks indicate genes that are listed in Table S1. (G) Heat-map indicates the major transcription factors that are important for HAM development and function, with similar pattern in AML cells and HAM. (H-J) STRING protein-protein interaction analysis of three key signaling pathways in HAM (PPARG, TGFB1, and CSF2). Most of the interacting proteins in these pathways are shown in white, indicating that they have similar expression levels in HAM and AML cells.

Next, we assessed transcription factors that are important for AM development and function. Of the PPAR family, PPAR-γ is critical for AM development (7) and was significantly upregulated in AML cells and HAM **(Fig. 4G)**, which corresponds with increased protein levels in AML cells **(Fig. 1R)**. RXRB associates with PPAR-γ, regulates cell differentiation, lipid metabolism and immune function, and was also highly expressed in HAM. The transcription factor KLF4, along with PPAR-γ, upregulates MCL-1 expression, and both were increased in HAM and AML cells. Downstream of GM-CSF (CSF2), PU.1 (SPI1) was increased in HAM and AML cells, both at the gene **(Fig. 1J; Fig. 4G)** and protein level **(Fig. 1R)**. CSF2RB is the receptor for GM-CSF, an important transcription factor for macrophage development and surfactant catabolism (29). CEBPB, RUNX-1, TGFBR2, IRF6 and IRF7 are important transcription factor-associated genes and showed similar expression patterns in AML cells and HAM relative to MDM **(Fig. 4G)**. They were highly expressed in HAM and associated with their development and function. We further performed STRING protein analysis on PPARG, TGFB1, and GM-CSF (CSF2), which mediate key signaling pathways in HAM. Most of the interacting proteins in these pathways showed similar expression in HAM and AML cells, indicating that these cells activate similar signaling networks **(Fig. 4H-J)**.

Thus, AML cells showed a similar transcriptomic landscape to HAM. Although MDM showed some similarity in the gene profile with HAM, especially in regard to common inflammatory and immune function-related pathways, important differences in canonical pathways exist (17). In addition, the expression level of major transcription factors important for AM development differed between MDM and AML cells and HAM **(Fig. 4G)**. Ingenuity pathway analysis (IPA) showed involvement of the RXRA transcription factor with upregulation of MARCO, COLEC12, HBEGF, IGF1, S100A4, and VCAN in AML compared to the MDM transcriptome **(Fig. S4B)**. Similar to a previous finding in HAM (17), the TREM1 signaling network was also upregulated in the AML transcriptome when compared to MDM **(Fig. S4C)**. IPA also identified the inflammatory response network as being distinct between AML cells and MDM, with involvement of PPARG and down regulation of CD36 in AML cells compared to MDM, a profile more consistent with HAM **(Fig. S4D)**. IPA network analysis identified network 1 (immune cell trafficking, cellular movement, cell-to-cell signaling and interaction), network 2 (cellular movement, immune cell trafficking, inflammatory response), and network 3 (immune cell trafficking, cellular movement, hematological system development and function) as being distinct between AML cells and MDM **(Fig. S4E-G)**. In summary, although MDM showed some similarity in the gene profile with HAM, AML cells have a transcriptomic profile more closely aligned with HAM.

### AML cells have upregulated lipid uptake genes and a drive towards oxidative phosphorylation

An important function of AMs is to regulate lipid metabolism, including degradation of lipid-rich surfactant to maintain proper lung function (30, 31). The lung alveolar space is rich in surfactant proteins and lipids, and has low levels of glucose (32, 33), an environment that may be conducive to the known low immunoreactivity of AMs, representing an adaptation to their specific environment. AMs engage in OxPhos over glycolysis as their core source of ATP (34), a finding in both human and murine cellular studies. Patients suffering from sepsis endure a shift from OxPhos to aerobic glycolysis, which is reversed upon patient recovery (35). *In vitro* studies established that LPS-stimulated inflammatory macrophages typically depend on glycolysis and alternatively activated M2 macrophages use OxPhos to generate energy (36). It is also well-known that IL-10 suppresses glycolysis in LPS-activated wild-type bone-marrow-derived macrophages (BMDMs) (37). Conversion from OxPhos to glycolysis in macrophages is generally important in host defense (38).

We assessed the metabolic status of AML cells relative to HAM and MDM. RNA-seq data demonstrated that expression of OxPhos-related genes is upregulated in AML cells and HAM relative to MDM **(Fig. 5A)**. Cholesterol and triglyceride metabolism-related genes are also upregulated in AML cells and HAMs **(Fig. 5B)**. Further, like reported for HAM (33, 34), we observed that AML cells exhibited much higher basal and maximal oxygen consumption rate (39)(OCR) compared with MDM, signifying the engagement of OxPhos and mitochondrial activities **(Fig. 5C).** Notably, AML cells also had a higher basal, maximal, and spare respiratory capacity (SRC) **(Fig. 5D-F)**. Proton leak, and non-mitochondrial oxygen consumption rates were elevated in AML cells compared to MDM **(Fig. 5G-I)**. These data support our RNA-seq analysis, where OxPhos is upregulated in HAM and AML cells **(Fig. 5A, B)**. Interestingly, the opposite trends were noted for extracellular acidification rate (ECAR) in AML cells, which indicate that the glycolytic rate (glycoPER), basal proton efflux rate, and compensatory glycolysis are much lower in AML cells as compared to MDM **(Fig. 5J-M)**. The rate of oxygen consumption can be increased for the production of ATP through either glycolysis or OxPhos (40). The ATP rate assay indicated that the production of ATP is increased in AML cells by OxPhos relative to MDM **(Fig. 5H)**. MDM are more prone towards glycolytic-linked ATP production than AML cells or HAM **(Fig. 5J).** We further assessed the metabolic response of cells after LPS stimulation, which was correlated with the glycolytic response. We measured extracellular lactate release in the supernatant after 24h of post treatment in AML cells and MDM. In contrast to MDM, AML cells did not respond to LPS to induce glycolysis **(Fig. 5N)**, similar to a previous finding reported in human AMs (34). Mitochondrial mass, mitochondrial membrane potential, the rate of proton leak, oxygen consumption, and ATP synthesis dynamically influence mitochondrial ROS production (41). We observed that mitochondrial-ROS (mt-ROS) and non-mitochondrial-ROS are increased in AML cells, as demonstrated by confocal microscopy and flow cytometry **(Fig. 5O-R)**. The data were further validated by measuring Electron paramagnetic resonance (EPR)-based mitochondrial ROS detection, which demonstrated an increase in the Mito-TEMPO**·** signal intensity in AML cell lysate **(Fig. 5S, T)**. Overall, these data provide evidence that AML cells are driven towards OxPhos rather than glycolysis and are consistent with the RNA-seq data indicating that fatty acid metabolism is more active in HAM and AML cells when compared to MDM.

**Fig 5.**
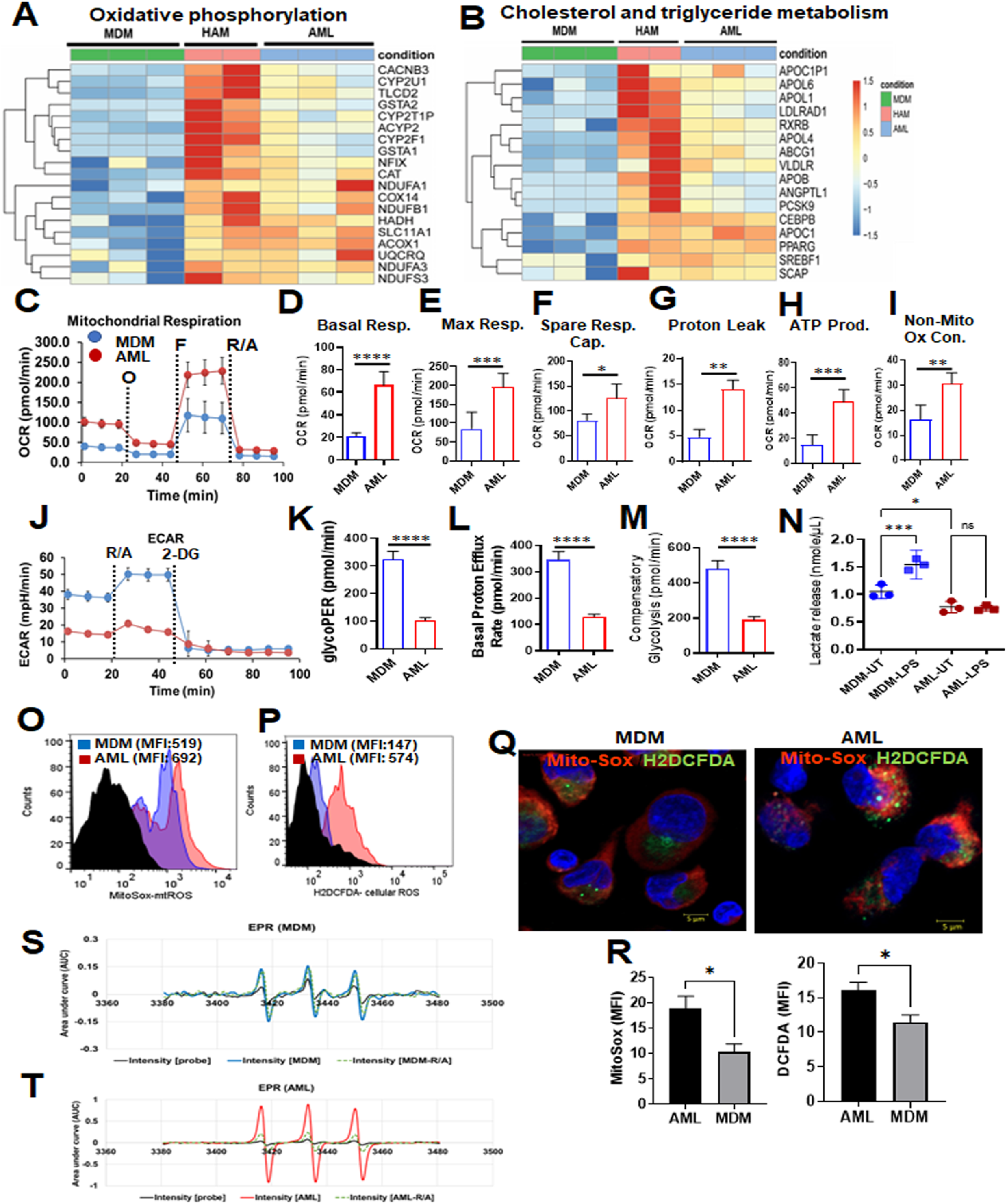
Metabolic status of AML cells, HAM and MDM. (A) Heatmap from the RNA-seq data indicates higher relative expression of genes related to fatty acid oxidation and Ox-Phos in AML cells and HAM. (B) Heatmap from the RNA-seq data indicates that cholesterol and triglyceride metabolism-related genes have a similar expression pattern in HAM and AML cells. (C-M) Red bars and lines represent AML cells, blue bars and lines represent MDM. Extracellular flux analysis performed in AML cells and MDM cells by Seahorse analyzer. The oxygen consumption rate (OCR) and extracellular acidification rate (ECAR) were analyzed under basal conditions and in response to Mito Stress Test reagent. (C) The dashed lines indicate when O: oligomycin; F: FCCP; R/A: rotenone and antimycin A were added. (D-F) Representative Mito Stress test kinetic graphs show higher levels of basal, maximal OCR and higher Spare respiratory capacity (SRC) in AML cells compared to MDM. (G-I) Proton leak, non-mitochondrial OCR and ATP production were also higher in AML cells. (J) The glycolytic rate (ECAR) kinetics graph demonstrates an increase in glycolytic rate in MDM as compared to AML cells. 2-deoxy-D-glucose (2-DG) was used to inhibit glycolysis. (K-M) Quantification of basal and compensatory glycolysis in MDM and AML cells. Representative experiment is shown of n=3, mean ± SD and analyzed by Unpaired Student’s ‘t’ test *p≤0.05, ** p≤0.01, ***p≤0.001, ****p≤0.0001. (N) Lactate levels (nonomol/µL) in the culture supernatant of MDM and AML cells after 24h LPS treatment (MDM: 10 ng/mL and AML: 100 ng/mL) was measured by Lactate Colorimetric Assay Kit II. Each dot represents an individual donor (n=3), mean ± SEM and analyzed with one-way ANOVA. *p≤0.05, ***p≤0.001. (O-Q) AML cells and MDM were treated with MitoSox (5µM) and DCFDA (5µM) to demonstrate mitochondrial and cellular ROS (non-mitochondrial), respectively, by flow cytometry and confocal microscopy. Magnification: 63x, Scale bar: 5µM. (R) Bar graphs show mitochondrial and cellular ROS represented as MFI. Representative experiment is shown of n= 3, mean ± SD and analyzed by Unpaired Student’s ‘t’ test *p≤0.05. (S, T) Electron paramagnetic resonance (EPR) spectrum-based mitochondrial ROS detection in MDM (blue line) and AML cells (red line) probed with Mito-TEMPO-H for signal intensity measurements in cell lysates. The data were analyzed first after baseline correction and subsequently second integration that yielded the area under curve (AUC) in arbitrary units (AU).

### Phenotypic and functional characterization of AML cells

HAM differentially express several cell surface receptors such as CD206, CD64, CD11c, CD163, CD170, MARCO, HLA-DR, CD11b and CD36 which can be used to distinguish HAM from other cell types (42-44). We characterized AML cell surface receptors by flow cytometry and confocal microscopy. We found that the AML cells exhibit increased expression of CD64, CD206, MARCO, CD163, CD11c, MerTK and CD170, and reduced expression of CD11b, CD36 and HLA-DR compared to MDM **(Fig. 6A-J; Fig. S5A)**. These data were verified by confocal fluorescence microscopy, which revealed higher expression of CD200R, CD11c, CD206, CD163, MARCO, MerTK, CD170, CD68 and CD64 in AML cells, and down regulation of CD36 and CD11b **(Fig. 6K-P; Fig. S5B-C)**. These changes in protein levels corresponded with the qRT-PCR data, where MRC1 and MARCO were highly expressed in AML cells *vs.* MDM, and CD36 was highly expressed in MDM *vs.* AML cells **(Fig. 1C, D, M).** MARCO is a scavenger receptor which mediates binding and ingestion of unopsonized environmental particles (44). MARCO is highly expressed in AML cells compared to MDM **(Fig. 1D; 6C, O, P)** and AML cells have higher capacity to bind unopsonized fluorescent beads compared to MDM **(Fig. S5D, E)**.

**Fig 6.**
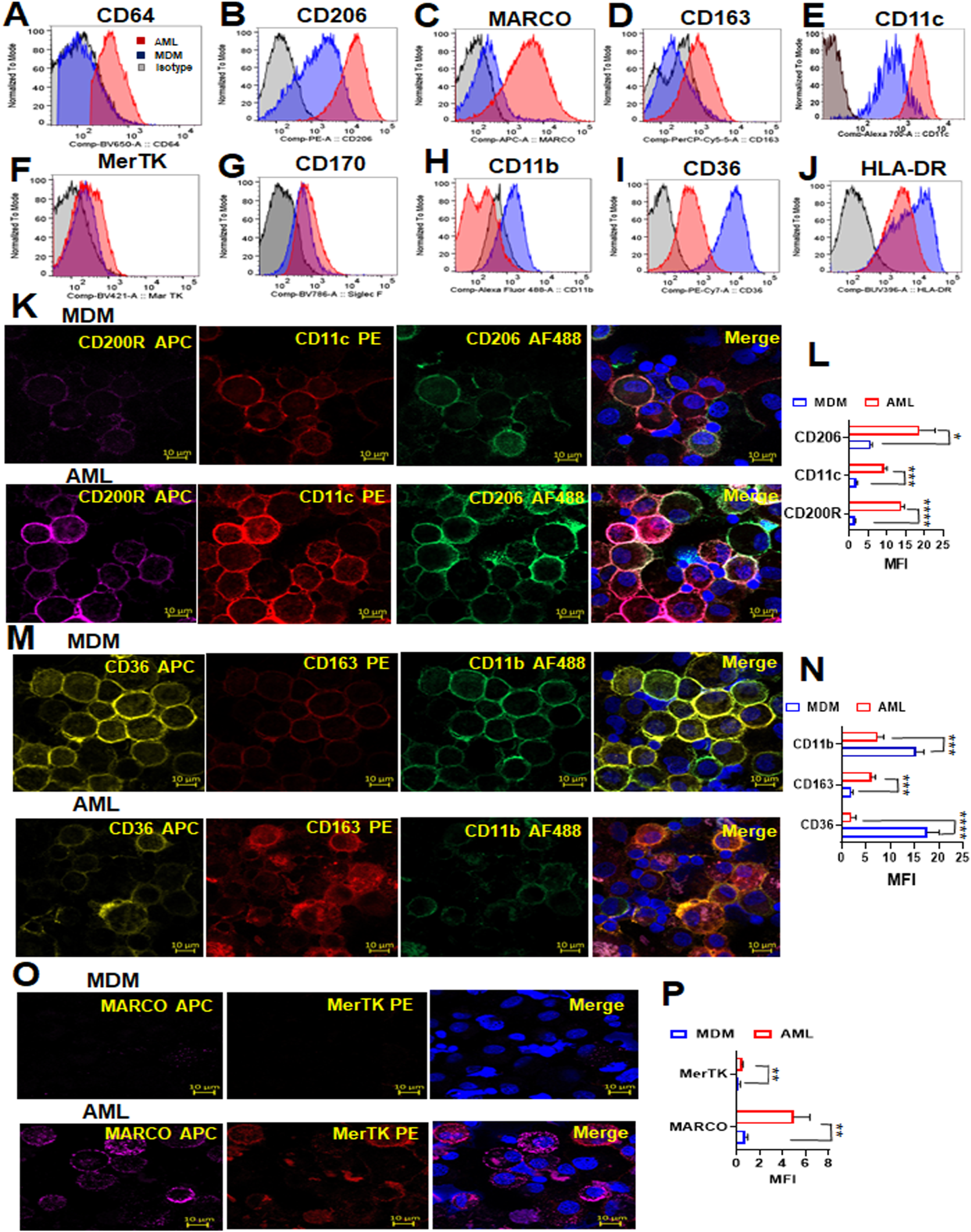
Phenotypic and functional characterization of AML cells compared to MDM. (A-J) PBMCs were exposed to ALL cocktail for 6 days on alternative days or left untreated (MDM). Flow cytometry data reveal that the AML cell surface phenotype resembles HAM with increased expression of (A) CD64, (B) CD206, (C) MARCO, (D) CD163, (E) CD11c, (F) MerTk and (G) CD170, and decreased expression of (H) CD11b, (I) CD36 and (J) HLA-DR when compared to MDM. Control fluorescence is shown in grey, and specific fluorescence for AML cells in red and MDM in blue. (K, M, O) The cells were immunostained with the indicated antibodies and DAPI for nucleus (blue), then imaged with confocal microscopy. Scale bar: 10µm and 63x magnification. (L, N, P) Confocal data were quantified by MFI and represented as bar graphs. Representative experiment of n = 3, mean ± SD and analyzed by Unpaired Student’s ‘t’ test *p≤0.05, ** p≤0.01, ***p≤0.001, ****p≤0.0001.

### AML cells secrete inflammation-related proteins

We assessed the amounts of secreted inflammation-related proteins from AML cells compared to MDM. Several secreted proteins reported for HAM were present in higher amounts (CD163, CXCL18, IL-13 and IL4) in AML cells compared to MDM **(Fig. 7A-D)**. Also similar to the profile reported for HAM, the levels of MMP7, MMP9, CCL22, TNFα and IFNγ were decreased significantly in AML cell supernatants compared to MDM **(Fig. 7E-I)**, which correlates with the RNA-seq data **(Fig. 4F).** Soluble ICAM-1, MCSF, IFNα, RAGE and IL1β levels in AML cell supernatants were similar to MDM **(Fig. 7J-N)**. We found GM-CSF and IL-10 in the cell supernatants as would be expected **(Fig. 7O, P)**. Similar profiles were observed in AML cells differentiated from purified monocytes **(Fig. 7Q-V)**.

**Fig 7.**
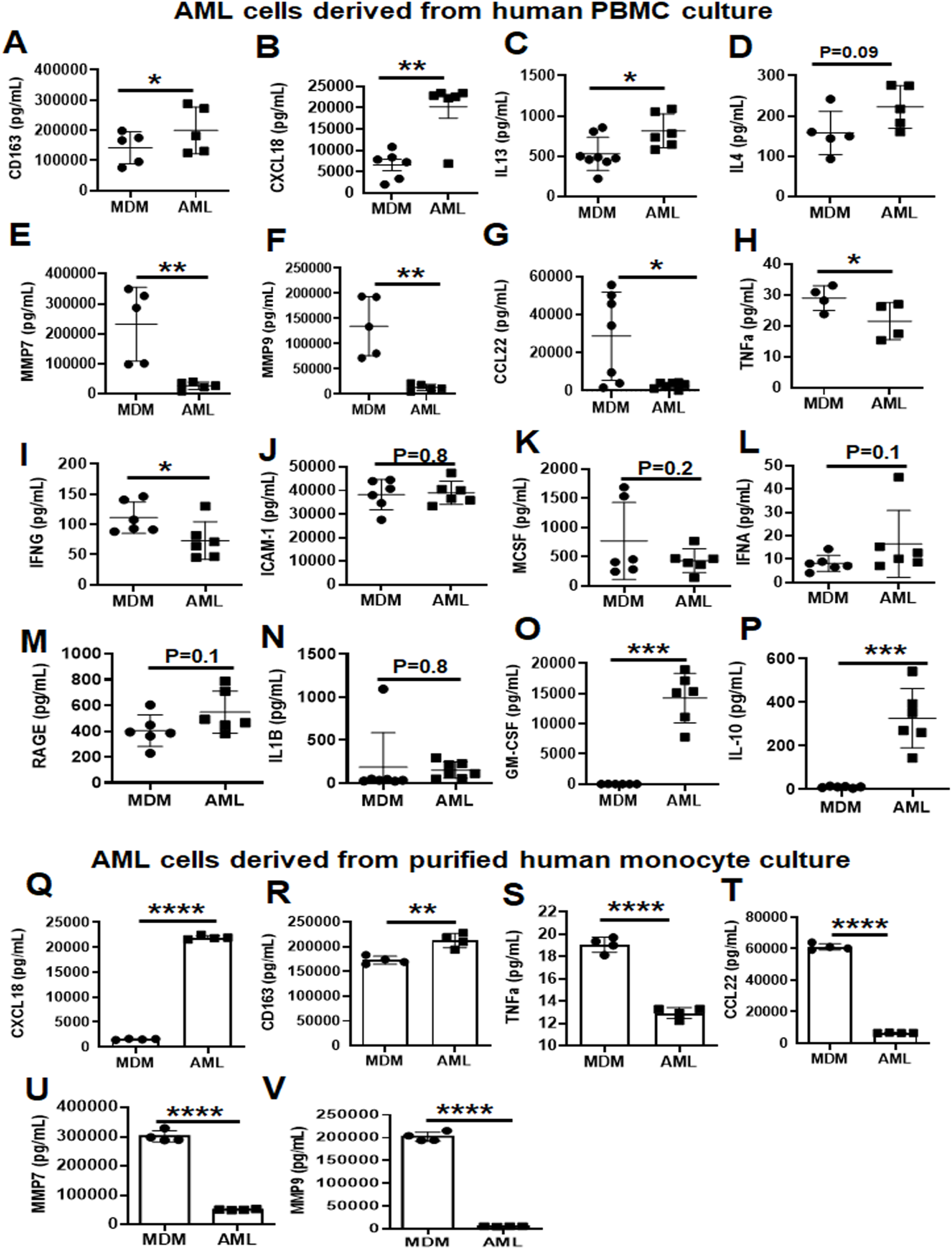
AML cells release several inflammation-related proteins. (A-I) PBMCs were exposed to ALL cocktail [surfactant (Infasurf) and cytokines (GM-CSF, TGF-β, IL-10)] for 6 days on alternative days or left untreated (MDM). Cell supernatants were collected and release of several inflammation-related proteins was analyzed simultaneously by Luminex technology. Like HAM, AML cells released increased levels of (A) CD163, (B) CXCL18, (C) IL13 and (D) IL4, and decreased levels of (E) MMP7, (F) MMP9, (G) CCL22, (H) TNFα and (I) IFNG compared to MDM. AML cells and MDM released similar quantities of soluble (J) ICAM-1, (K) MCSF, (L) IFNA, (M) RAGE and (N) IL1B, and there was significantly more (O) GM-CSF and (P) IL-10 in the supernatants collected from AML cells than from MDM. Data are mean ± SEM; each dot indicates results from 1 donor (n=5-8) and analyzed by Unpaired Student’s ‘t’ test. *p≤0.05, ** p≤0.01, *** p≤0.001. (Q-V) Monocytes were purified by EasySep™ human monocyte isolation kit by negative selection magnetic sorting of healthy human PBMCs on Day 0 and exposed to ALL cocktail treatment [surfactant (Infasurf) and cytokines (GM-CSF, TGF-β, IL-10)] for 6 days on alternative days or left untreated (MDM). Cell supernatants were collected and release of inflammation-related proteins was analyzed simultaneously by Luminex technology. AML cells differentiated from isolated monocytes release higher (Q) CXCL18 and (R) CD163, and lower (S) TNFα, (T) CCL22, (U) MMP7 and (V) MMP9 amounts than MDM, similar to those cells differentiated from PBMCs. Data are expressed as mean ± SEM; each dot indicates results from 1 donor (n=4) and analysed by Unpaired Student’s ‘t’ test. ** p≤0.01, ****p≤0.0001.

### HAM and AML cells demonstrate increased uptake and intracellular growth of *M.tb,* and increased persistence of SARS-CoV-2 compared to MDM

AMs are the first myeloid cells to phagocytose airborne *M.tb* and allow for *M.tb* growth (45-47). To investigate the phagocytic capacity of HAM, AML and MDM, we infected each cell type with mCherry-H_37_R_v_ *M.tb*. We found that HAM and AML cells have similar, increased phagocytic capacity compared to MDM as demonstrated by confocal microscopy **(Fig. 8A, B).** In addition to calculating mean bacteria per cell, we found that 62.6% ± 1.5 HAM and 59.8% ± 1.5 of AML cells contained ≥ 1 bacterium, compared to 41.6% ± 8.4 of MDM (mean ± SEM, n = 3-4). We also found increased intracellular *M.tb* growth in HAM and AML cells over time **(Fig. 8C)**. Similarly, murine AMs are more permissive to *M.tb* infection and growth than BMDMs (16).

**Fig 8.**
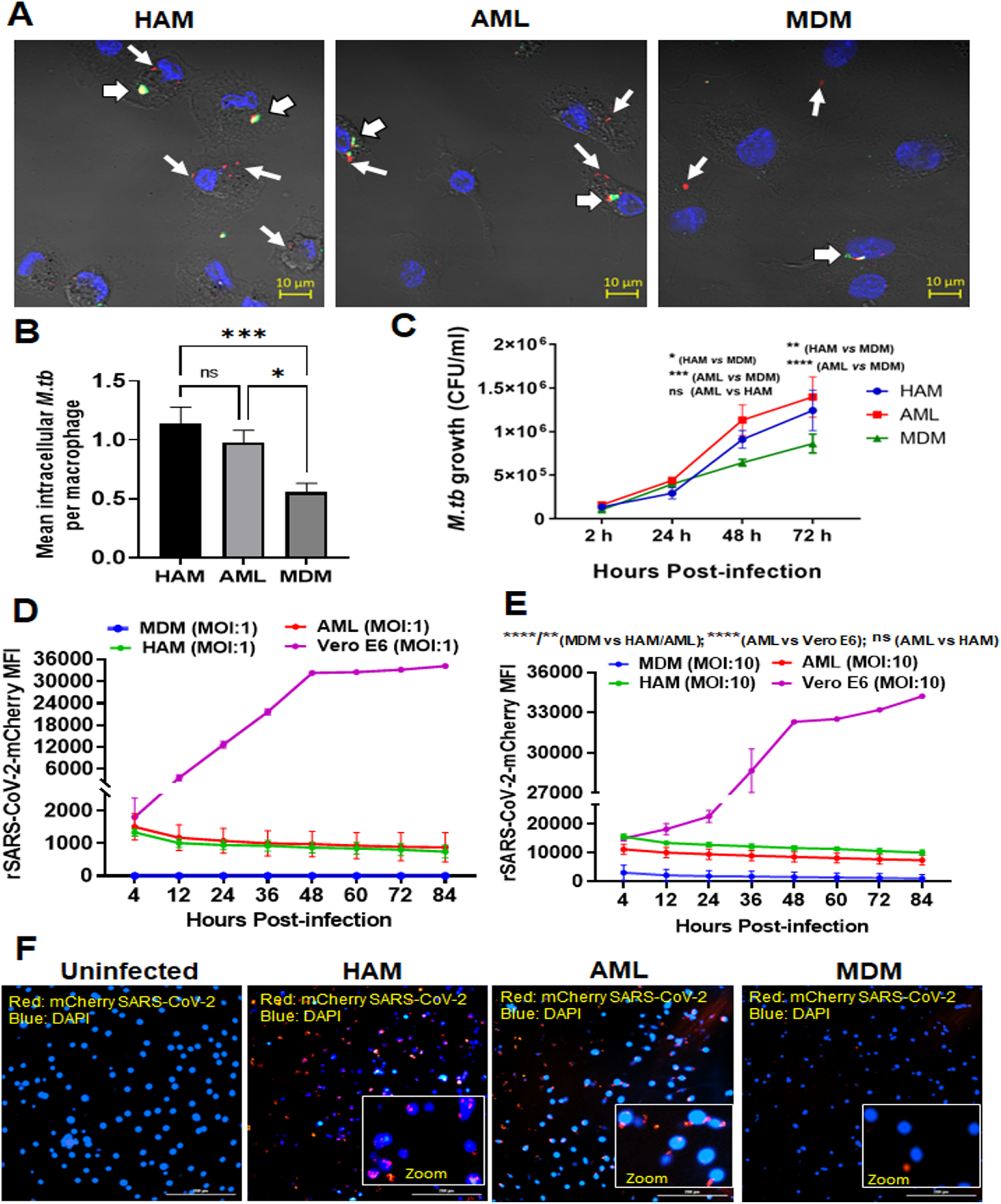
Uptake and growth of *M.tb* and SARS-CoV-2 in AML cells are similar to HAM. PBMCs were exposed to ALL cocktail for 6 days on alternative days or left untreated (MDM). Freshly isolated HAM were obtained from the same donor. Cell monolayers were then incubated with *M.tb*-H_37_R_v_-mCherry (MOI 5; red) for 2h, fixed without permeabilization and washed. (A) Cell monolayers on coverslips were immunostained with fluorophore-conjugated anti-*M.tb* antibody (green) and DAPI (blue), and then imaged using confocal microscopy. 63x magnification, Scale bar: 10 µm. White arrows indicate mCherry (Red) intracellular *M.tb* and white arrowheads indicate attached/extracellular (Yellow-green) *M.tb.* (B) Mean number of intracellular bacteria per cell was calculated from >100 macrophages. A representative experiment from MDM/AML cells n = 5, HAM n=3, mean ± SD. Data were analyzed by Ordinary one-way ANOVA. * p ≤ 0.05, *** p ≤ 0.001. (C) Intracellular growth of *M.tb*-H_37_R_v_ was monitored in the indicated time points post-infection (2, 24, 48, 72h) by CFUs. Each point is the mean of CFU values from triplicate wells. Representative experiment of n=5, mean ± SD with 2-way ANOVA. * p ≤ 0.05, ** p ≤ 0.01; *** p ≤ 0.001; ****p = <0.0001. (D, E) Kinetics of increased uptake of SARS-CoV-2 (MOI:1 and 10) and persistence over time using the Cytation 5 live cell imaging system. Data were normalized to uninfected control and presented as mCherry MFI values. Data are expressed as mean ± SD with one-way ANOVA. ** p ≤ 0.01; ****p = <0.0001. (F) Representative image of mCherry positive cells infected with SARS-CoV-2/mCherry-Nluc, counter stained with DAPI at day 5 post infection. Red: mCherry SARS-CoV-2, blue: DAPI (nucleus). Scale bar: 200 µm and 20x magnification. Insert photomicrographs show higher power images of cells infected with rSARS-CoV-2/mCherry-Nluc (Red). The data in D-F are representative of 4 experiments using different MDM/AML donors and a HAM donor. Videos of cells infected with rSARS-CoV-2/mCherry-Nluc using Cytation 5 live cell imaging 4-84h post-infection are shown in Movies S1-S4.

We next explored the cellular response to SARS-CoV-2 infection. The Vero E6 kidney cell line is extensively used in COVID-19 research for viral propagation, passaging and stock preparation, and antiviral assays (48). These cells highly express the ACE2 receptor for SARS-CoV-2 attachment, but lack the co-receptor TMPRSS2 protease that also participates for entry into human cells (49) Viral entry into Vero E6 cells is reported to be cathepsin-mediated but may not mimic viral infection of human cells (50). Cells expressing both ACE2 and TMPRSS2 are highly permissive for infection. We confirmed that Vero E6 cells express ACE2 receptor but not TMPRSS2. ACE2 basal level receptor expression was higher in AML cells than MDM but lower than Vero E6 cells **(Fig. S6A)**. TMPRSS2 basal level receptor expression was higher in AML cells than MDM. Vero E6 cells did not express TMPRSS2 **(Fig. S6B)**. Basigin (CD147) is another reported route of cellular entry for SARS-CoV-2 (51). We found that BSG/CD147 expression is similar in AML cells, MDM and Vero E6 cells **(Fig. S6C)**.

The cellular response to SARS-CoV-2 in primary human macrophages, particularly HAM, has not been explored. We infected HAM, AML cells, MDM and VeroE6 cells using our previously described replication-competent recombinant SARS-CoV-2 expressing bioluminescent luciferase (Nluc) and fluorescent mCherry reporter genes (rSARS-CoV-2/mCherry-Nluc) (52) and continuously monitored viral infection over time by the Cytation 5 fluorescence live cell imaging system **(Fig. S6D; Movies S1-S4)**. These data demonstrated that similar to HAM, AML cells have rapid, increased SARS-CoV-2 uptake and persistence over time without replication in contrast to Vero cells, where SARS-CoV-2 entry is initially lower but propagation is higher **(Fig. 8D, E; Movies S1-S4)**. Finally, we assessed the SARS-CoV-2 persistence after DAPI counterstaining in HAM and AML cells at day 5-post infection **(Fig. 8F)**. We observed lower infection and persistence of SARS-CoV-2 in MDM than HAM or AML cells.

## Discussion

We developed an AML cell model using blood-derived monocytes that recapitulates unique features of AMs, which require TGF-β and GM-CSF for development along with IL-10 for maintenance and have a critical role in catabolizing lipid surfactant. We show that culturing human blood-derived monocytes, either purified or with other PBMCs, in a cocktail containing surfactant, IL-10, TGF-β and GM-CSF allows monocytes to differentiate into AML cells and maintain this phenotype over time in culture. Using multiple complementary approaches, we demonstrate that these AML cells mimic AMs in that they have 1) similar morphology containing lipid bodies; 2) similar gene expression patterns with only 6.4% of detectable genes showing a significant change in expression between AML cells and HAM; 3) increases in gene and protein levels of CD206, PPAR-γ, MARCO and other key markers for AMs; 4) similar expression of genes in key pathways required for AM development (PPAR-γ, TGF-β and GM-CSF); 5) specific histone modifications with higher levels of H3K4me1 and lower levels of H3K4me3; and 6) increased OxPhos and reduced glycolysis. Importantly, AML cells demonstrate increased uptake and intracellular growth and persistence of *M.tb* and SARS-CoV-2, respectively, similar to HAM. Both AML and MDM cells demonstrated very limited proliferation capacity as has been described for healthy adult HAMs (27, 28). In contrast to MDM, AML cells did not respond to LPS to induce glycolysis, as reported for HAM (34). Although, previous data also suggest that virulent *M.tb* can drive the shift towards aerobic glycolysis in HAM (39). These findings suggest the differential metabolic activity of HAM in response to a pathogen.

Study of HAM has been hindered by the invasive and costly nature of BALs, which require extensive prescreening tests and experienced pulmonologists. In contrast, venipuncture is significantly cheaper, less invasive, and requires less specialized training than needed for the BAL procedure. People are generally more willing to undergo venipuncture than BAL, and venipuncture can occur more frequently, thus making recruitment of donors easier for studying blood-derived cells. Our AML model is based on culturing blood monocytes with commercially available products, and AML cells can be purified from the lymphocytes through a simple and inexpensive adherence step, thus providing a HAM model that is much more readily available to a range of labs.

A second challenge in working with HAM is that each BAL yields approximately 2-4 x 10^6^ HAM per person, thus restricting studies to smaller experiments than what is feasible when working with cell lines. In contrast, from a full blood draw, approximately 50 x 10^6^ monocytes (and thus potentially AML cells) can be recovered per donor, allowing for much larger studies. Cell number is a particular problem when working with mice because BAL results in only approximately 1-2 x 10^5^ AMs per mouse. Thus, many investigators pool cells from multiple mice to have sufficient cells for one experiment. This need makes studying heterogeneity in the population challenging. The ability to work with blood-derived AML cells obviates the need to pool samples, allowing for studies that interrogate donor heterogeneity. Finally, some important inflammatory pathways differ in humans and mice (53). Thus, working with primary human cells has a distinct advantage. Studies using non-human primate (NHP) BAL cells to obtain AMs is another option. However, there is more limited access to NHPs, studies are expensive and there is strict IRB regulation similar to humans. It will be interesting to adopt our AML model to NHP monocytes for greater accessibility and application.

An alternative approach to studying AMs is to digest animal, typically murine lungs, and study total lung macrophages. This results in the recovery of many more macrophages than BAL. However, the lung contains a range of macrophages, including alveolar, interstitial, and intravascular. AMs constitute about 10% of lung macrophages and have a unique phenotype relative to interstitial and intravascular macrophages; thus, study of lung macrophages is not specific to AMs and the majority of recovered macrophages from lung tissue are actually IMs. AMs can be isolated from digested lung tissues based on cell surface receptor expression, but the cell yield is still low.

A third challenge with studying AMs is that their phenotype rapidly changes after removal from the lung, which makes long-term studies and mechanistic work challenging. Importantly, we show that culturing AML cells in Infasurf, IL-10, TGF-β and GM-CSF allows them to better retain a HAM phenotype once adhered. This is expected to allow investigators to conduct longer mechanistic studies than what is currently feasible *in vitro*, and so represents a large step forward for the field.

Several studies have highlighted the unique susceptibility of HAM to airborne bacterial and viral infections (12, 45, 46, 54). In an effort to begin to demonstrate the applicability of AML cells for infection studies, we analyzed the uptake and intracellular growth of *M.tb* and SAR-CoV-2 and determined that these parameters were similar in AML cells and HAM. For example, AML cells had increased phagocytosis and intracellular growth of *M.tb* (47). Regarding SARS-CoV-2, we detected moderate expression of ACE2 and TMPRSS2 in AML cells. It is of interest that AML cells demonstrated rapid uptake of SARS-CoV-2 and subsequent persistence without detectable growth. Viral persistence of SARS-CoV-2 is reported in research and clinical settings (55, 56). Infection of NHPs showed that rhesus macaques and baboons develop moderate SARS-CoV-2 viremia with COVID-19-related pathology and some degree of viral persistence (57). Analysis of human PBMCs showed viral persistence in the form of fragmented SARS-CoV-2 RNA and the presence of S1 viral proteins in the post-acute sequelae of COVID-19 (PASC) patients for up to 15 months post-acute infection (58). Finally, a humanized mouse model identified that SARS-CoV-2-infected human lung resident macrophages contribute to hyperinflammation of the lung by upregulation and release of IL-1, IL-18 and activation of the inflammasome (59). AML cells hold promise for further investigation of the molecular and cellular events enabling SARS-CoV-2 uptake and persistence in human cells.

Prior studies have generated macrophages from primary human cells using specific factors such as GM-CSF or MCSF to generate either M1 or M2 types of macrophages. HAM shares both M1 and M2 characteristics and thus cannot be characterized as such (42). Our approach was to optimize dosage in combining critical factors into a cocktail (Infasurf, GM-CSF, TGF-β, IL-10) in order to better recapitulate the alveolar environment of AMs. We based the concentration range of factors to study on prior in vitro studies. It is difficult to measure the true concentration of these factors in the alveolar hypophase. We did not include all known soluble factors of the alveolar hypophase (e.g., eicosanoids) in our model but found that factors previously found to be critical for AM development recapitulated the AM phenotype well. In future study, it will be interesting to explore the role of other factors such as eicosanoids (esp. PGE_2_) in AML longevity (26).

As noted above, previous studies, including our own, have determined that AMs will change their phenotype when removed from the lung (16, 17). Thus, isolation and handling of AMs could affect their transcriptome and raise the concern that the ex vivo-studied AMs have deviated from the in vivo phenotype. Studying human AMs in vivo is not possible. We contend that isolated HAM analyzed soon after harvest (within 6h) without manipulation most closely approximates the in vivo phenotype (17). Including the cocktail during in vitro cultivation maintains this phenotype. The inclusion of these factors needs to be considered when studying these cells in a variety of lung inflammation contexts. Finally, although we demonstrate that cultivating human monocytes in the cocktail enables cells to differentiate in a manner that more closely recapitulates HAM, use of MDM in culture has generated very significant and useful data over the years in delineating human macrophage innate immune responses confirmed in vivo and in humans. Thus, the success of this and other new models does not necessarily diminish the importance of data from older models.

A bottleneck in studying respiratory biology has been the ability to study AMs *in vitro*. The lungs are a unique organ that must balance pro- and anti-inflammatory responses and much of this is mediated through AMs, which have a unique biology relative to other tissue macrophages. We expect that this AML model will significantly advance respiratory biology research, including for inflammatory diseases like chronic obstructive pulmonary disease (COPD), asthma, and cystic fibrosis, and infectious diseases including COVID-19 and tuberculosis. This model should also aid in assessing the impact of aging on AM biology. Finally, this model will aid in translational human studies. i.e., therapies and vaccines. Thus, we expect this AML model will help in identifying key pathways/responses to interrogate in the context of the lung alveoli and can be further validated in less readily available primary HAM as necessary.

## Materials and methods

### Ethics Statement

PBMCs were isolated from human peripheral blood collected from healthy adult donors, following a Texas Biomed approved IRB protocol (20170315HU). HAM were isolated from BAL of healthy adult human donors, following a Texas Biomed approved IRB protocol (20170667HU). All donors for these studies provided informed, written consent.

### Collection and isolation of HAM

Fresh HAM were isolated and cultured from BAL of healthy donors as described previously (17) and used for the respective studies. See the detailed supplemental methods section.

### MDM culture

PBMCs were isolated from individual adult healthy donors by Ficoll-Paque cushion centrifugation using an established protocol (60). The cells were then cultured in sterile Teflon wells (2×10^6^/ml) with RPMI 1640 +10% fresh autologous serum at 37°C/5% CO_2_ for 6 days to allow for differentiation of monocytes into MDM (60, 61).

### *In vitro* development of AML cells and MDM from human PBMCs

PBMCs were cultured for 6 days to allow for differentiation of monocytes into untreated MDM or treated AML cells. To generate AMLs, Infasurf (100 μg/ml), GM-CSF (10 ng/ml), TGF-β (5 ng/ml) and IL-10 (5 ng/ml) were added on days 0, 2, and 4 (three doses of ALL cocktail). For some experiments, Infasurf, GM-CSF, TGF-β, and IL-10 were only added on day 0 (one dose of ALL cocktail). In other experiments, we analyzed the role of individual components of the ALL cocktail. On day 6, both control MDM and AML cells were harvested and adhered to tissue culture dishes for 2 h in RPMI 1640 with 10% fresh autologous serum, lymphocytes were washed away, and then all experiments were performed. We also determined the requirement of the continuous addition of ALL cocktail after differentiation to retain the AML cell phenotype. The complete protocol and surfactant components information are in detailed supplemental methods section.

### Isolation of human monocytes by magnetic sorting and development of AML cells

PBMCs were obtained for CD14 positive monocyte isolation using the EasySep Human Monocyte Isolation Kit (Stem cell Technologies), according to the manufacturer’s instructions. Isolated monocytes were cultured for 6 days to allow for differentiation of purified monocytes into AML cells or untreated MDM. AML and MDM cell lysates and supernatants were used for qRT-PCR and Luminex assay, respectively, to compare differentiated AML cells from PBMCs or freshly isolated monocytes. See more information in the detailed supplemental methods section.

### Cytospin analysis

Single cell suspensions of freshly isolated HAM, cultured AML or MDM cells (5×10^4^) were placed in a cytofunnel and centrifuged at 150 x g for 5 min onto cytoslides that were dried and stained with HEMA 3 differential staining. The slides were examined with an AE2000 microscope. See details in the detailed supplemental methods section.

### Transmission electron microscopy (TEM)

AML and MDM cells were fixed with 4% formaldehyde and 1% glutaraldehyde in phosphate buffer (Invitrogen) overnight at 4°C. The samples were processed and imaged using a JEOL 1400 TEM. Expanded protocol is in the detailed supplemental methods section.

### Ki67 cells proliferation assay

MDM, AML and THP-1 monocytic cells (3×10^5^ cells/polystyrene FACS tubes) were collected, then fixed and permeabilized by adding 300 µl 100% methanol (pre-stored at −20 ◦C) for 5 min at RT. Cells were then washed by centrifugation (250g for 10 min) with FACS cells staining buffer (Cat# 420201, BioLegend). Cells were next treated with human 5µl/tube TruStain FcX™ (Fc receptor blocking solution, Cat# 422302, BioLegend) and incubated for 30 min at RT. The cells were then stained with 5µl/tube Alexa Fluor 488 anti-Ki67 antibody (BD Bio Science, Cat# 561165) and respective Alexa Fluor 488 Mouse IgG1 k isotype matched control for 45 min at 4◦C. Cells were then washed with cells staining buffer by centrifugation. Flow cytometry samples (∼2×10^5^/tube) were analyzed using a BD FACS symphony instrument and the data were analyzed using Flowjo software.

1×10^5^ cells were placed in a cytofunnel and centrifuged at 150 x g for 5 min onto cytoslides. The coverslips were placed with mounting reagent ProLong Gold Antifade Mountant with DAPI (Thermo Fisher Scientific). The cells on slides were visualized with a Ziess LSM 800 confocal microscope (20X and 63X magnification) and counted based on DAPI staining using Image J FiJi software. The percentage of Ki67 positive cells was calculated from >200 macrophages (DAPI positive cells) per microscopic field.

### RNA isolation, quantification and qRT-PCR

Cultured AML and MDM cells were harvested and RNA isolated using the manufacturer’s RNA extraction protocol (Invitrogen). cDNA was prepared. Real time PCR was performed using predesigned TaqMan human primers in the Applied Biosystems 7500 Real-Time PCR System. Expression levels of basal mRNA in AML and MDM cells were normalized to ACTB and calculated by the ΔΔ threshold cycle (ΔΔCT) method. The detailed protocol is in the detailed supplemental methods section.

### Multi-color flow cytometry

Single cell suspensions of AML and MDM cells were incubated with fluorochrome tagged antibodies along with their respective isotype matched control antibodies. Samples were analyzed using a multi-color BD FACS symphony instrument and the data were analyzed using Flowjo software. The expanded description and gating strategy used is presented in **Fig. S5** in the detailed supplemental methods section.

### Multicolor confocal microscopy

AML and MDM cells were stained with fluorescence-conjugated antibodies or control antibodies and stained slides were visualized with a Ziess LSM 800 confocal microscope. See in the detailed supplemental methods section.

### Bead cell association study

AML and MDM cells were incubated with non-opsonized FluoSpheres™ Sulfate Microspheres, 1.0 µm, yellow-green fluorescent F8852-beads (Invitrogen). Stained slides were visualized with a Ziess LSM 800 confocal microscope. The cells were counted based on DAPI staining. The number of fluorescent beads was also counted and shown as beads/macrophage. See further information in the detailed supplemental methods section.

### RNA-Seq and analyses

Freshy isolated HAM and 2h adherent MDM and AML cells were lysed in TRIzol and RNA was isolated using Direct-zol RNA Miniprep kit, R2052 (Zymo Research) as per the manufacturer’s instructions. RNA sequencing was carried out using the HiSeq 3000 platform (Illumina). The detailed protocol and data analysis are elaborated in the detailed supplemental methods section.

### Luminex-multiplex analysis

Luminex assays were performed in the culture supernatants of AML and MDM cells following the manufacture’s protocol by the Luminex® 100/200™ System. The analytes IL-1β, IL-2, IL-4, IL-6, IL-8, IL-10, IL-12 p70, IL-13, IL-18, TNFα, IFN-α, IFN-γ, CCL5, CCL8, CCL18, CCL22, CD163, GM-CSF, ICAM-1, M-CSF, MIF, MMP-1,

MMP-7, MMP-9, MMP-12 and RAGE were detected by Luminex Human Discovery Assay. The data were analyzed by Belysa™ Immunoassay Curve Fitting Software (Millipore Sigma). See information in the detailed supplemental methods section.

### Western blot analysis

AML and MDM cells were lysed with NE-PER Nuclear and Cytoplasmic Extraction kit according to the manufacturer’s protocol (Thermo Scientific). Western blot was performed and the membranes were incubated with the primary antibody for PPAR-γ, PU.1, H3K4me1 and H3K4me3 followed by anti-rabbit IgG, HRP-linked antibody. The membranes were developed using clarity ECL reagent on a UVP chemstudio 815 system. Stripping was performed and probed to detect β-actin or Histone H3 levels. See further information in the detailed supplemental methods section.

### Extracellular Flux Analysis

Real-time cellular metabolism of AML cells and MDM was determined by measuring OCR (pmol/min) and ECAR (mPh/min) using a Seahorse Analyzer, according to the manufacturer’s instructions (Agilent Technologies). Mito stress assay was performed after sequential addition of 5 μM oligomycin, 4 μM FCCP and 2 μM rotenone and antimycin A. For glycolysis stress analysis, AML and MDM cells were injected with 2 μM rotenone and 2 μM antimycin A followed by 100 mM 2-deoxyglucose (2-DG) to determine glycolytic rate. See information in the detailed supplemental methods section.

### Lactate release

MDM and AML cells were stimulated with or without LPS (MDM: 10 ng/ml and AML: 100 ng/ml) for 24h. The supernatants (collected from 3 donors and stored at −80◦C) were used for lactate measurements. Lactate was quantified using a lactate colorimetric enzymatic assay kit according to the manufacturer’s instruction (K627, BioVision Inc, Milpitas). Data were expressed as per manufacturer’s instruction nmole/µl cell culture supernatant.

### Mitosox assay and cellular ROS detection

Mitochondrial and intracellular non-mitochondrial ROS levels in AML and MDM cells were measured by staining with Mitosox (5μM) and H2DCF (5μM), respectively, for confocal or flow cytometry analysis. See further information in the detailed supplemental methods section.

### EPR assay

Electron paramagnetic resonance (EPR)-based ROS detection for mtO_2_ was performed using Mito-Tempo-H (100 µM). 2 μM rotenone and 2 μM antimycin A mix were added to inhibit mitochondrial complexes. The EPR spectra were measured on the Bruker EMXnano ESR system. Expanded methods are in the detailed supplemental methods section.

### Phagocytosis assay for *M.tb*

Fixed HAM, AML or MDM monolayers were incubated with either rabbit polyclonal anti-*M.tb* whole cell lysate antibody or an IgG rabbit isotype control antibody. Later, cells were incubated with an AlexaFluor 488 donkey anti-rabbit secondary antibody. Imaging was executed on a Zeiss LSM 800 microscope. The complete protocol and analysis are in the detailed supplemental methods section.

### Macrophage infection with *M.tb*

MDM, HAM and AML cells were infected (MOI 2) with virulent *M.tb* H_37_R_v_. *M.tb* intracellular growth at each post-infection time point (2, 24, 48 and 72h) was measured in cell lysates. CFUs were assessed after 3 and 4 weeks on 7H11 agar plates. Expanded information are in the detailed supplemental methods section.

### Generation of rSARS-CoV-2 expressing reporter genes

Recombinant SARS-CoV-2 expressing mCherry and nanoluciferase (Nluc) reporter genes (rSARS-CoV-2/mCherry-Nluc) was rescued in Vero E6 cells and viral stocks prepared (52). Viral titers in the stocks were determined and used for infection studies. Complete information is in the detailed supplemental methods section.

### Macrophage infection with rSARS-CoV-2/mCherry-Nluc

HAM, AML, MDM and Vero E6 cells were infected with rSARS-CoV-2-mCherry-Nluc virus (MOI: 1 or 10 PFU/cells). The infected cells were used for further study (see detailed supplemental methods section).

### Cytation 5 live cell imaging assay

Freshly obtained HAM, AML, MDM and Vero E6 cells were infected with rSARS-CoV-2/mCherry-Nluc virus (MOI: 1 and 10). Live cell imaging was performed using Citation 5 paired with BioSpa (Biotek/Agilent). Data analysis was achieved with Gen5 software by calculating mCherry MFI. Cells were counted after 120h by counter staining with DAPI. Persistence of SARS-CoV-2-mCherry in cells was monitored in time lapse videos (0/4-84h time period) using Gen5 software. The extended protocol is in the detailed supplemental methods section.

### Statistical analyses

Graphs were prepared and statistical comparisons applied using GraphPad Prism version 9 (GraphPad). Statistical comparisons were performed by unpaired two-tailed Student’s t-test. Ordinary one-way and two-way analysis of variance (ANOVA) with Sidak’s multiple comparisons test for multiple-testing (GraphPad Prism 9) was applied wherever applicable (indicated in the figure legends). For Correlation analysis, Spearman’s rank test was applied. Statistical differences between groups were reported significant when the p-value was ≤ 0.05. The data are presented as mean ± SEM.

## Data Availability Statement

RNA-seq data can be found in the NCBI GEO database: https://www.ncbi.nlm.nih.gov/geo/query/acc.cgi?acc=GSE188945.

## Author Contributions

SP and LSS designed these studies. SP, EA, JS, AA, IG, ZL, NJ, ML and CY conducted the experiments and acquired the data. SP, EA, JS, AA, YW, HZ, HC, DJM, JP, JBT and LMS analyzed the data and performed critical review; SP, EA, LSS wrote the paper.

## Acknowledgement

This work was supported by NIH award [AI136831] (to LSS), Texas Biomed Cowles and Forum Postdoctoral Fellowship (to SP) and NIH award [1R01AI161175-01A1] and [1R01AI161363-01] (to LMS). Research reported in this publication was supported by the NIH-NIAID under Award Number P30AI168439. Research reported in this publication was also supported by the Office of The Director, NIH Award [S10OD028653]. RNA sequencing data were generated in the Genome Sequencing Facility, supported by UT Health San Antonio, NIH-NCI P30 CA054174, NIH Shared Instrument grant [1S10OD021805-01] (S10 grant), and CPRIT Core Facility Award [RP160732]. TEM was conducted at the Electron Microscopy Laboratory at UT Health San Antonio and fluorescence microscopy imaging was conducted with instruments at the Biology Core at Texas Biomed. We thank Dr. Colwyn A. Headley for his guidance with the seahorse metabolism experiments.

## Supplemental materials

### Supplemental detailed methods

Additional detailed methods are provided along with references.

### Supplemental Figures

**Fig. S1. Selection of the optimal dose of GM-CSF, TGF-β and IL-10 and role of lung-associated components individually and in combination in generating AML cells.** PBMCs from healthy human donors were exposed to lung-associated component treatment [cytokines GM-CSF (2.5, 5, and 10 ng/mL), TGF-β (2.5, 5, and 10 ng/mL) and IL-10 (2.5, 5, and 10 ng/mL)] for 6 days on alternative days or left untreated (MDM). The optimum concentration was selected by measuring gene expression of (A) PPARG and (B) MRC1 by qRT-PCR. Gene expression was normalized to Actin. Representative bar diagram showing the relative mRNA expression expressed as mean ± SD (n=2 donors). PBMCs from healthy human donors were exposed to lung-associated component treatment [surfactant (Infasurf: 100 µg/mL) and cytokines (GM-CSF: 10 ng/mL, TGF-β: 5 ng/mL, IL-10: 5 ng/mL) alone or all together (ALL treated)] for 6 days on alternative days (Day 0, 2, 4), or left untreated (MDM). qRT-PCR data demonstrated significant increases in expression of (C) PPARG, (D) MARCO, (E) CCL18, (F) MRC1, (G) CES1 and (H) MCEMP1, and decreases in (I) MMP9, (J) CD36 and (K) MMP7 in the ALL cocktail group compared to treatment with each component alone. The data were normalized to the actin control. Data are expressed as mean ± SD (n=2). Data are expressed as mean ± SD and analyzed with one-way ANOVA * p≤0.05, ** p≤0.01, ***p≤0.001, ****p≤0.0001. (n=2). (L) ALL cocktail treatment does not affect the viability of AML cells. PBMCs were exposed to Infasurf (100 µg/mL) and cytokines (GM-CSF: 10 ng/mL, TGF-β: 5 ng/mL, IL-10: 5 ng/mL) (ALL cocktail treated)] for 6 days on alternative days (Day 0, 2, 4), or left untreated (MDM). AML cells were untreated or treated with 0.1 % triton X-100 for 5 min to induce cell and nuclear membrane breakage (positive control). Cells were then stained with Annexin V (ANXA5/annexin V-APC) followed by Ethidium Homodimer-2 (EthD-2, 4µM) to assess for cell viability. The numbers in the insets indicate the percentage of Annexin V and EthD-2-positive cells. Representative data from 3 donors.

**Figure S2. Continuous supplementation of the lung-associated components is required to retain the HAM phenotype of AML cells.** PBMCs were exposed to lung-associated components treatment [Infasurf (100 µg/mL) and cytokines (GM-CSF: 10 ng/mL, TGF-β: 5 ng/mL, IL-10: 5 ng/mL)] for 6 days on alternative days, or left untreated (MDM). The adherent macrophages were then treated with or without ALL of lung-associated components [Infasurf (100 µg/mL) and cytokines (GM-CSF: 10 ng/mL, TGF-β: 5 ng/mL, IL-10: 5 ng/mL)] and incubated for 24, 48, and 72h. At each time point cells were collected to quantify the expression of (A) PPARG, (B) MRC1, and (C) MARCO by qRT-PCR. Blue circle dots show individual donors of the MDM control for 0h (day 6), 24h, 48h or 72h (without any treatment). Green triangles indicate individual donors of the AML cells treated with 3 doses of cocktail up to day 6 and cultured for 0h (day 6), 24h, 48h or 72h without additional treatment. Red squares indicate individual donors of AML cells where additional treatment supplementation (Post treated) after adherence can extend the HAM phenotype. n=3, Mean ± SEM.

**Figure S3. Continuous supplementation of the lung-associated components retains the HAM phenotype of AML cells for longer duration.** Monocytes were purified from human PBMCs by magnetic sorting and cultured with lung-associated components [Infasurf (100 µg/mL) and cytokines (GM-CSF: 10 ng/mL, TGF-β: 5 ng/mL, IL-10: 5 ng/mL)] for 6 days on alternative days, or left untreated (MDM). On day 6, the differentiated cells were plated (5×10^5^/ well) in a 24 well plate. Adherent cells were then treated with or without ALL cocktail (post treatment) on days 2, 4 and 6 (see illustration). At each time point cells were collected to quantify the expression of (A) PPARG, (B) MRC1, and (C) MARCO by qRT-PCR. Blue circle dots show individual donors of the MDM control for day 0 (day 6 of culture), 2, 4 or 6 (without any treatment). Green triangles indicate individual donors of the AML cells treated with 3 doses of cocktail up to day 6 differentiated culture and then cultured for day 0, 2, 4, or 6 without additional cocktail treatment. Red squares indicate individual donors of AML cells treated with 3 doses of ALL cocktail up to day 6 differentiated culture and then cultured for day 0, 2, 4 or 6 with additional treatment supplementation (Post treated) on day 0, 2 and 4. n=2, Mean ± SEM.

**Fig. S4. AML cells and MDM share transcriptional signatures and related pathways.** (A) Volcano plot demonstrates the comparison between the AML and MDM transcriptome. AML cells and MDM are more dissimilar than AML cells and HAM: out of 14,097 expressed genes, 744 are up-regulated ≥2-fold with FDR adjusted p-value < 0.05 (red), and 438 are down-regulated ≥2-fold with FDR adjusted p-value < 0.05 (blue) in MDM. (B-D) IPA analysis identified several pathways containing genes that were significantly up-regulated in AML cells relative to MDM with similar expression in AML cells and HAM. They include involvement of (B) RXRA transcription factor with upregulation of MARCO, COLEC12, HBEGF, IGF1, S100A4 and VCAN, (C) TREM1 and (D) Inflammatory response network with involvement of PPARG and down regulation of CD36. (E-G) IPA network analysis of genes that were differentially expressed in AML cells compared to MDM identifies (E) network 1 (immune cell trafficking, cellular movement, cell-to-cell signaling and interaction), (F) network 2 (cellular movement, immune cell trafficking, inflammatory response) and (G) network 3 (immune cell trafficking, cellular movement, hematological system development and function).

**Fig. S5. Flow cytometry gating strategy for Figure 6 A-J, confocal and cell association assays.** (A) FSC vs SSC was used as the initial gate, then FSC vs FSC to gate singlets. The population was then gated on CD64 positive cells (macrophages) for subsequent analysis. (B) Cells were immunostained with the indicated antibodies and DAPI for nucleus (blue), then imaged with confocal microscopy. AML cells had higher CD170, CD68 and CD64 expression than MDM. Scale bar: 10µm and 63X magnification. (C) Confocal data were quantified by mean fluorescence intensity (MFI) and are represented as a bar graph. Representative data showing the expression of indicated proteins of 3 different donors. Data are expressed as mean ± SD and analyzed by Unpaired Student’s ‘t’ test ** p≤0.01, ****p≤0.0001. (D) Cell association study using unopsonized green fluorescent beads comparing AML cells and MDM, nuclei were stained with DAPI (blue). Scale bar: 10µm and 63X magnification. (E) Quantification of the number of beads per macrophage. Representative experiment is shown of n=2, Mean ± SD, **** p < 0.0001.

**Fig. S6. AML cells express ACE2, TMPRSS2, and BSG/CD147 and have increased uptake and persistence of SARS-Cov-2 monitored by Cytation 5 live cell imaging.** (A-C) ACE2, TMPRSS2 and BSG/CD147 receptor expression was measured in MDM, AML cells and Vero E6 cells by qRT-PCR. n = 3 human donors. (D) mCherry fluorescent SARS-CoV-2 in individual wells was monitored for uptake and persistence over time by the Cytation 5 live cell imaging system (MDM, HAM, AML cells and Vero E6 cells). Representative mini-graphs (of n=4) demonstrate total mCherry fluorescence units (Y-axis) versus post infection time (X-axis). Shown up to 84h post infection.

**Movie S1. Video of SARS-CoV-2 mCherry viral infection in MDM using Cytation 5 live cell imaging 0/4-84h post-infection.** Persistence of SARS-CoV-2-mCherry virus over time was monitored by Cytation 5 live cell imaging. The time-lapse video shows mCherry (red) SARS-CoV-2.

**Movies_MPEG4 files\MDM_SARS-CoV-2_MOI 1.mp4**

**Movie S2. Video of SARS-CoV-2 mCherry viral infection in AML cells using Cytation 5 live cell imaging 0/4-84h post-infection.** Persistence of SARS-CoV-2-mCherry virus over time was monitored by Cytation 5 live cell imaging. The time-lapse video shows mCherry (red) SARS-CoV-2.

**Movies_MPEG4 files\AML_SARS-CoV-2_MOI 1.mp4**

**Movie S3. Video of SARS-CoV-2 mCherry viral infection in HAM using Cytation 5 live cell imaging 0/4-84h post-infection.** Persistence of SARS-CoV-2-mCherry virus over time was monitored by Cytation 5 live cell imaging. The time-lapse video shows mCherry (red) SARS-CoV-2.

**Movies_MPEG4 files\HAM_SARS-CoV-2_MOI 1.mp4**

**Movie S4. Video of SARS-CoV-2 mCherry viral infection in Vero E6 cells using Cytation 5 live cell imaging 0/4-84h post-infection.** Persistence of SARS-CoV-2-mCherry virus over time was monitored by Cytation 5 live cell imaging. The time-lapse video shows mCherry (red) SARS-CoV-2.

**Movies_MPEG4 files\VeroE6_SARS-CoV-2_MOI 1.mp4**

